# Gas modulating microcapsules for spatiotemporal control of hypoxia

**DOI:** 10.1101/2022.09.02.506302

**Authors:** Thomas G. Molley, Shouyuan Jiang, Chantal Kopecky, Chavinya D. Ranaweera, Gagan K. Jalandhra, Jelena Rnjak-Kovacina, Kristopher A. Kilian

## Abstract

Oxygen is a vital molecule involved in regulating development, homeostasis, and disease. The oxygen levels in tissue vary from 1 to 8% with deviations having major biological consequences. In this work, we developed an approach to encapsulate enzymes and nanozymes, at an unprecedented loading capacity, which precisely controls the oxygen content in cell culture. Here, a single microcapsule is able to locally perturb the oxygen balance, and varying the concentration and distribution of matrix embedded microcapsules provides spatiotemporal control. We demonstrate attenuation of hypoxia signaling in populations of stem cells, cancer cells, endothelial cells, and cancer spheroids. Capsule containing hydrogel films applied to chick chorioallantoic membranes encourage neovascularization, providing scope for topical treatments or hydrogel wound dressings. We further demonstrate versatility by loading capsules with ceria nanorods as “nanozymes” to modulate active oxygen species with potential as a cytoprotective treatment. The approach can be used in multiple formats, including deposition in hydrogels, as granular solids for 3D bioprinting, and as injectable biomaterials. Overall, this platform’s simplicity and flexibility will prove useful for fundamental studies of oxygen-mediated processes in virtually any *in vitro* or *in vivo* format, with scope for inclusion in biomedical materials where controlling hypoxia may be clinically advantageous.

## 1. Introduction

Oxygen is one of the most important molecules that drives life on earth. For humans and other mammals, oxygen tension plays a critical role in aiding tissue development and maintaining normal tissue function, as well as pathological processes like tumor progression.^1^ Low levels of oxygen in tissue is termed hypoxia, and hypoxic conditions mediate the transcriptional activity of hypoxia inducible factors (HIF) which regulate gene expression associated with multiple functional activities including angiogenesis, cell migration, differentiation, metabolism, and apoptosis.^2,3^ These HIF proteins are sensitive to the local oxygen tension which varies greatly within and between different tissues of the body.^1,4^ While air has a saturation of around 20% oxygen, circulating blood levels sit around 5%,^3^ with various tissues such as bone marrow and cartilage reaching as low as 1%. ^5^ These values contrast significantly from the 18.6% oxygen found in cell culture incubators,^6^ leading to cellular hyperoxia *in vitro*. For decades, creating an easy and effective way to fine tune the oxygen levels for *in vitro* cultures has remained a challenge.

The gold standard for mimicking low oxygen environments *in vitro* is the use of hypoxic incubators and hypoxic chambers. These devices control atmospheric oxygen levels and thus limit the amount of diffusible oxygen within the cell culture environment. However, there is little granularity with these tools. They are expensive, inaccessible to many researchers, and their oxygen levels reset when opened.^7,8^ To overcome these issues, researchers have explored the use of chemical and enzymatic methods to create hypoxic environments. Baumann et al. first explored this concept in 2008 by using glucose oxidase paired with catalase to induce hypoxia in three different cancer lines.^9^ Further work also explored the use of laccase as an oxygen depletion enzyme to reduce the consumption of glucose from cell media.^10^ Critically, free enzymes are not stable long term in cell culture as they can readily be degraded by proteases. Therefore, to gain further control over hypoxia in hydrogels, Gerecht and colleagues synthesized hydrogels with laccase covalently conjugated to the polymer backbone to aid in enzyme stability.^11^ This technique has also been reported by Dawes et al. using glucose oxidase immobilized hydrogels.^12^

While these systems enable greater control of local oxygen levels, the methodology is specialized and limits the selection of the hydrogel network. Therefore, to broaden this concept’s versatility, we sought an approach that could apply to any hydrogel system while utilizing the benefits afforded via enzymatic oxygen control. To this end, we aimed to entrap the enzymes within microcapsules. Since the substrates of glucose oxidases and catalase are small molecules, a polymer membrane that is large enough to enable small molecule diffusion, but too large to accommodate enzyme diffusion is needed. The capsules would require gentle processing such that a high quantity of active enzymes could be integrated, and the presence of capsules should not disrupt bulk network properties of the hydrogel or resident cells. Currently, there are no microencapsulation approaches that satisfy these criteria.

In this paper, we demonstrate the first microscale gas-modulating system for spatiotemporal control over oxygen tension in cell culture. The novel microencapsulation system uses a gelatin microgel template with a polydopamine coating, where tuning the coating polymerization alters the permeability and stability of capsules. Our approach led to unprecedented concentrations of enzyme, up to 40 mg/mL within the microcapsules, with stable activity maintained for over a month in cell culture conditions. We observed a doseresponse in oxygen levels with enzyme concentration and demonstrated functional outcomes through reproducible control over stem cell and cancer cell activities. Local attenuation of oxygen levels was shown to increase neovascularization in 2D, 3D, and *in ovo* models, providing scope for these materials to be used in promoting in *vivo* angiogenesis. As a demonstration of versatility, nanozymes were added to capsules as an alternative approach to modulate the gas balance, demonstrating reduction in active oxygen species to protect adipose derived stem cells in a tissue irradiation model. Overall, this work demonstrates a versatile system where gas-modulating capsules can be integrated with virtually any soft material system to locally control gas levels.

## 2. Materials and methods

### 2.1 Gelatin microcapsules synthesis

For empty capsules, type B Gelatin (Bloom strength 300) was dissolved at 10 wt% in 1X PBS in a 40 °C water bath (75 mL of oil). For loaded capsules, a 20 wt% solution of gelatin was dissolved, and it was mixed 1:1 with a solution containing the cargo. The gelatin solution (2 mL) was pushed through a 0.45 μm filter into an oil bath (olive oil) under constant stirring at 38 °C. After 10 minutes of equilibration, the oil bath was cooled to 10 °C, using a surrounding ice bath, and left to stabilize for 30 minutes. Hexane (25 mL) was subsequently added and left to stir for 10 minutes before the microgels were decanted off into centrifuge tubes. The particles were left to settle for 30 minutes prior to 4 washing steps with hexane. The particles were agitated in the hexane solution and quickly pipetted onto a 25 μm nylon filter over Kim wipes in a fume hood to let the hexane dry off. A spatula was used to spread out the particles on the filter to ensure all hexane was evaporated. Once hexane free, the particles were placed into a centrifuge tube again and resuspended in a Tris-HCl buffer (0.1M pH 8.5, 10 mL). Dopamine hydrochloride was added (final concentration of 10 mg/mL) and the tube was covered with a Kim wipe to allow oxygen exchange. The tube was then placed on an orbital shaker at 300 rpm for the specified time. The particles were then washed with 1X PBS four times and stored until further use. For size characterization, uncoated particles were placed onto a glass slide and imaged under an optical microscope. One hundred particles were imaged across 7 images, and the diameters were measured using ImageJ.

### 2.2 Encapsulation efficiency and Maximal loading

To quantify encapsulation efficiency, the capsule synthesis procedure was followed (loaded with 10, 40, or 250 kDa FITC-dextran cargo (20 μg/mL)) until just before adding in the dopamine. The particles were then placed on the orbital shaker for 4 hours to simulate the time prior to a stable coating forming on the surface of the microgels. Microgels were then weighted out and dissolved at 37 °C in 1X PBS to extract the contents. The contents were then measured using a fluoresce spectrophotometer with a laser excitation at 488 nm (Cary eclipse). The emission at 517 nm was plugged into a line of best fit from a standard curve to calculate the concentration of Dextrans in solution. For maximal BSA loading, the same procedure was followed expect the gelatin particles were loading with 100 mg/mL of Bovine Serum Albumin (BSA) and 125 μg/mL of fluorescent FITC-labelled BSA. A separate standard curve for FITC-BSA was produced to create a line of best fit.

### 2.3 Cell Culture and Maintenance

All cell culture was performed using aseptic techniques inside a sterile class II biological safety cabinet. All cells were cultured inside an incubator at 37°C with a 5% CO_2_ and 100% humidity atmosphere. All cells used in this study were between the passages 2-13. The media used for each cell line is described in **Table S1**. Media changes were performed every 2 days. The adipose derived stem cells (ADSC), B16F10, MCF-7, MDA-MB-231 and A375-P cells were all passaged when a confluency of 80-90% was reached. Human umbilical vascular endothelial cells (HUVECs) were passaged at 70-80% confluency. For all cell lines, 0.25% Trypsin-EDTA was used for detachment.

### 2.4 Immunofluorescence Staining and Tissue Clearing

Clearing solutions were prepared as described previously with minor modifications.^13^ Cubic solution 2 was prepared by mixing 50 wt% sucrose, 25 wt% urea, 10 wt% triethanolamine with DI water at 55°C until also fully dissolved. Microgel suspensions were fixed using 4 wt% Paraformaldehyde (PFA) for 1-4 days at room temperature to ensure full penetration of PFA into thick constructs. The gels were then rinsed with 1X PBS followed by 3 1X PBS washes at 2-4-hour intervals. Afterwards, the gels were incubated in 0.5 wt% triton overnight before repeating the same wash procedure. The staining for all gels was performed using the dyes and antibodies in Table S2. The gels were washed with 1X PBS three times after both primary and secondary staining before the addition of the Cubic 2 clearing solution for 1-4 days. All confocal imaging was performed with a Zeiss LSM 800. A 10x objective with a 2.5 mm working distance was used to see deeper into the samples. Samples were coated with clearing 2 solutions throughout the duration for the imaging to prevent drying.

### 2.5 Glucose oxidase activity characterization

Glucose oxidase activity was characterizing by using the glucose oxidase activity assay kit according to the manufacturer’s protocol. Briefly, glucose oxidase was first dissolved to 20 mg/mL (2000 U/mL) in a 50mM sodium acetate buffer at a pH of 5.0. The enzyme was mixed 1:1 with a 20 wt% Gelatin solution to make microcapsules that were coated for 24 hours. Two other microcapsules were then synthesized: one with 20% as much glucose oxidase, and one with no glucose oxidase. The capsules were then place on 25 μm nylon filters, to remove excess water, and were weighed into an Eppendorf tube. The particles were then resuspended to 20 mg/mL in the GOx buffer solutions from the kit. Then, 50 μL of the capsule buffer solution was added to 4 different wells in 96 well plate, along 4 wells for blanks, and 5 wells for the standard curve. The plate was then placed into a plate reader (CLARIO Star Plus) set to 37 °C to warm up. Finally, 50 μl of the kit working solution was added to each well and measurements were taken every 3 minutes for 45 minutes total. Unused microcapsules were then stored in 4 °C for 1 month, with new measurements taken on days 2, 7, 14, and 21.

### 2.6 Seeding cells with glucose oxidase and catalase microcapsules

For fixed cell experiments, glucose oxidase capsule laden microgel suspensions were prepared the same method used in **2.15**. For cancer cell response, microgel suspensions were loaded with A375-P (1 × 10^6^ cells per mL) and cultured for 1 and 7 days. Cells were subsequently fixed and stained with Hoechst, Phalloidin, and HIF1-alpha antibody. The images were subsequently taken on a Zeiss LSM 800 confocal and imaging (100 μm z-stack with 2 μm slices). Aggregate size analyses were performed in ImageJ. Briefly, the actin cytoskeleton of aggregates was thresholded and masked and the Analyze particles plugin was used to calculate the average aggregate size per replicate.

For the ADSC cell response, microgel suspensions (medium, 10 wt%, 1% filler) were loaded with 2 vol% microcapsules (0, 10, 25, or 100 U/mL glucose oxidase with a 1:60 ratio of catalase) and 1 × 10^6^ ADSCs per mL. For images, 100 μm z-stacks with 2 μm slices were taken. To calculate mean grey value intensities, the phalloidin channel was used to create ROIs of cell boundaries, and then theses ROIs were applied to the HIF1-alpha channel to achieve the mean grey of hif1-alpha staining per replicate region. For cell counting, the nuclei were thresholding and counted via the analyze particle’s function, and HIF1alpha expressing cells were counted manually.

### 2.7 Irradiation protection study

Two sets of microcapsules were prepared; one with 5 mg/mL of ceria nanoparticles loaded (dissolved in DI water), and one with 6 mg/mL of catalase loaded (dissolved in 1X PBS). Three separate microgel suspensions were made using medium-sized 10 wt% GelMa particles, 0.5 wt% filler, and 0.05 wt% of LAP: one with 20 mg/mL of nanoceria capsules, one with 20 mg/mL of catalase capsules, and one with no capsules. ADSCs were loaded into each suspension at 1 × 10^6^ cells per mL, and the suspensions were UV crosslinked in plastic molds. The crosslinked suspensions were placed in 24 wells plates with 1 mL of media and placed in an incubator for 2 hours. Then, the media was replaced with fresh media containing either 0, 10, 100, or 1000 μM of H2O2 and cultured for 24 hours. The media was then removed, and the sample were washed once with 1X PBS. A 1X PBS solution of Calcein AM (2 μM) and Ethidium Homodimer-1(4 μM) was added (500 μL) to each well and the gels were incubated at 37 °C for 40 minutes. Afterwards, the samples were washed with 1X PBS, placed back in the incubator for 5 minutes, and washed again before imaged. Quantification was performed using the same method as described in the supplemental methods.

### 2.8 Statistical analysis

All whiskers in box plots are one standard deviation (s.d.). For particle size study, 100 different particles were used. Four replicates were used for the permeability study, and 3 replicates used per condition for the encapsulation efficiency study. Five replicates were used for the BSA capsule stability study, as well as the maximal BSA loading test. For 3D printed Line fidelities, 8 separate lines were used, with the diameter averaged across 6 different points along the line. Quadruplicates were used for the Glucose oxidase activity studies, and triplicates for the dissolved oxygen tests for the capsules. Triplicates were used for the live dead studies. Quadruplicates were used for all ADSC studies, triplicates were used for the A375 studies, and six replicates were used for the breast cancer studies. No replicates were performed for the western blots. Five replicates were used for all HUVEC studies, and 3 eggs were used for each replicate in the CAM assay. Triplicates were used in the MitoSOX study and the irradiation model test. Statistical significance was determined using a oneway ANOVA with Tukey’s Post Hoc HSD analysis. Differences were considered significant when P < 0.05. Statistical significance was highlighted in figures with the following convention: * = P < 0.05; ** = P < 0.01; *** = P < 0.001, **** = P < 0.001

## 3. Results and Discussion

### 3.1 Characterization of polydopamine coated gelatin microparticles

Our method of oxygen reduction relies upon the dual reaction of glucose oxidase paired with catalase. The glucose oxidase enzyme first oxidizes a glucose molecule, forming a gluconic acid and hydrogen peroxide molecule. Catalase in turn breaks two hydrogen peroxide down into two water molecules and an oxygen molecule. Together, this reaction reduces one dissolved oxygen molecule for every two glucose molecules (**Figure 1A**). Previous use of this enzyme pair in vitro has been shown effective at creating hypoxia in cell culture conditions, but the free enzymes had poor stability and lost function quickly over time.^14^ We hypothesized that microencapsulation of these two enzymes in a stable hydrogel environment would provide a simple yet effective method to prolong enzyme life and stability while maintaining efficacy.

**Figure 1.**
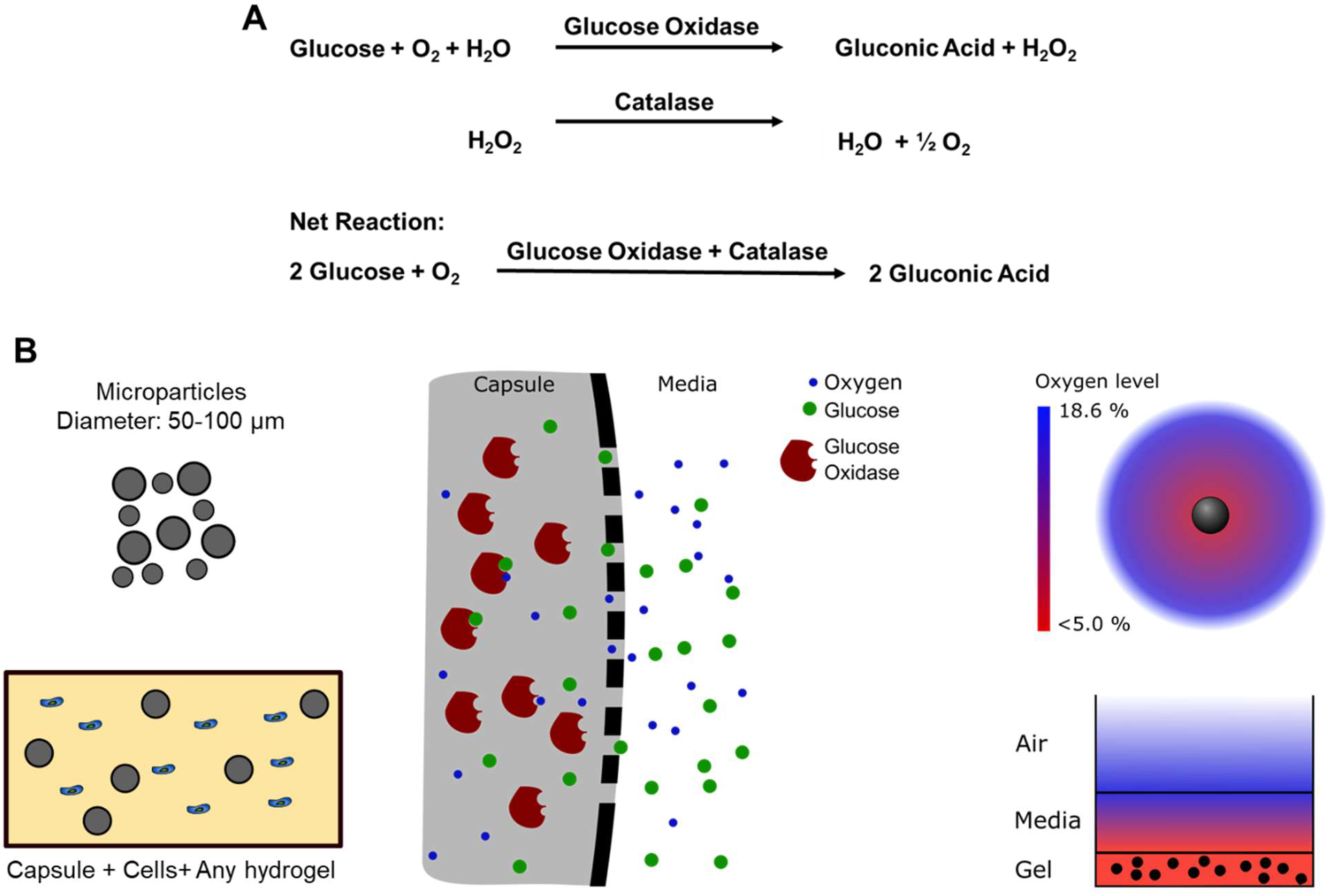
Schematic of hypoxic capsules in microgels. A) Reaction scheme of glucose oxidase and catalase. B) Representation of glucose oxidase enzyme within a capsule. The enzymes are entrapped within the capsule walls, but the wall pore sizes are small enough to enable the exchange or enzyme substrates (glucose and oxygen). (Right) Representation of oxygen levels around an individual capsules (top) and a capsule laden hydrogel in cell culture medium.

For the capsule design, gelatin was chosen as the capsule core due its low thermal gelation temperature which enables easy microgel formation while preserving payload bioactivity. Polydopamine (PDA) was chosen as the capsule wall since it coats most materials with ease and has a mesh size that is smaller than the enzymes but larger than glucose (Pore diameters: PDA ~ 1 nm; glucose oxidase radius ~ 6 nm).^15,16^ This enables diffusion of analytes in and out of the capsules while preventing enzymes from escaping or proteases from entering and degrading the enzymes. Once loaded, these capsules function as oxygen sinks, with ease of dispersion and spatial localization for control over oxygen tension (**Figure 1B**). The kinetics of oxygen reduction can subsequently be tuned by varying either the number of embedded capsules or the concentration of enzyme within each capsule. Spatial arrangement of capsules with different enzyme content is also possible to provide gradients of gas concentration.

The microcapsule synthesis scheme using a water in oil emulsion method is shown in Figure S2. To maximize loading potential, cargo is added to the gelatin solution at high concentration prior to addition to the oil bath. Importantly, previous methods to generate microgels of gelatin or gelatin methacryloyl (GelMa) with oil emulsions use acetone to dehydrate the particles prior to washing and collection.^17,18^ Here, acetone will render loaded enzymes inactive. Therefore, the particles require a non-water-miscible solvent to wash away excess oil. Hexane was chosen as it readily dissolves food oils while remaining immiscible with water.^19^ Optical images verify the formation of stable gelatin microgel particles without acetone dehydration (**Figure 2Ai**); scanning electron micrographs show smooth morphologies on the uncoated particle surfaces (**Figure 2Aii**).

**Figure 2.**
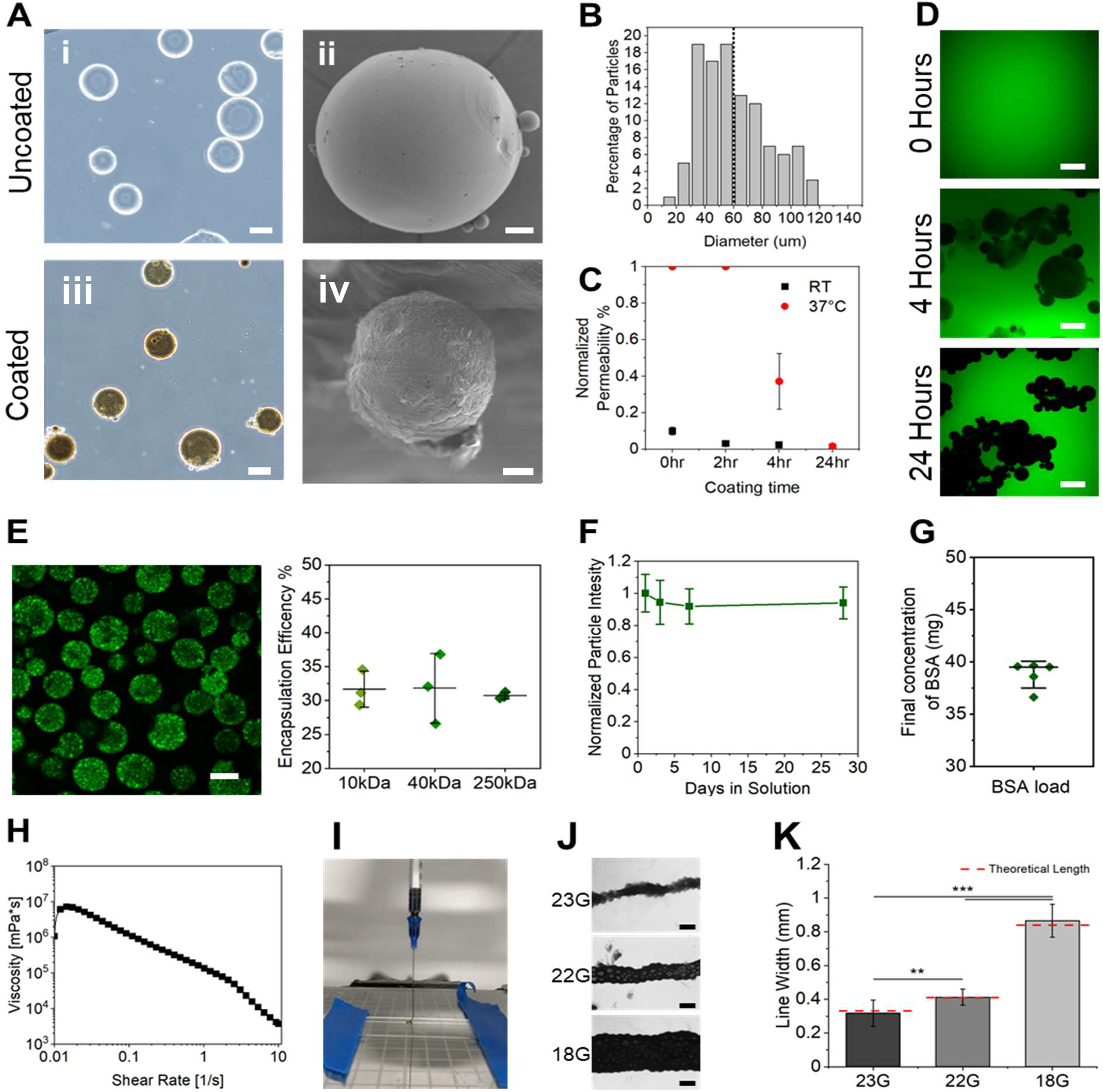
Characterization of PDA gelatin microcapsules. A) (i) Bright field microscopy images of uncoated gelatin microgels. Scanning electron microscope images of uncoated (ii) and PD coated (iv) gelatin capsules. (iii) Bright field microscopy image polydopamine (PD) coated gelatin microgels. B) Size distribution histogram of uncoated gelatin microgels. C) Normalized permeability of capsules based on fluorescence. D) Confocal microscopy images of capsules (coated for 0, 4, or 24 hours) sitting in FITC-Dextrans for 24 hours (37 °C). E) (Left) Confocal microscopy image of 24-hour PD coated microcapsules loaded with 250 kDa FITC-Dextran (2 mg/mL). (Right) Plot of encapsulation efficiencies calculated for 10, 40, and 20kDa dextrans. F) Plot of normalized fluorescence intensity of FITC-BSA in microcapsules after 28 days. G) Plot of the final concentration for 5 batches of BSA loaded microcapsules. H) Viscosity vs shear strain rate plot from shear strain sweeps (log ramp from 0.02-10 s^-1^). I) a photograph of a printed continuous line filament of jammed microcapsules. J) Bright field microscopy images of jammed microcapsule printed lines with various nozzle sizes (23, 22, and 18G). K) A plot of analyzed microcapsule line diameters with varied nozzle dimeters. Red lines indicate the inner diameter of the nozzles used. Scale bars: 300 μm. Scale bars: 10 μm (A (ii and iv)), 50 μm (A (i and iii), E), 300 μm (J). **P < 0.01, ***P < 0.001, ANOVA. Error bars represent s.d.

For the coating, dopamine hydrochloride (10 mg/mL) was added to a dilute suspension of gelatin microgels (10 w/v% particles in Tris-HCL Buffer (pH 8.5)) and shaken in a capless centrifuge tube to enable gas exchange. After 24 hours, the particles adopted a brown/black appearance (**Figure 2Aiii**), corresponding to the color of polydopamine (PD).^20,21^ SEM showed a slightly rougher surface on coated particles (**Figure 2Aiv**), and size characterization revealed an average particle size of 60 ± 24 μm (**Figure 2B**). To determine the barrier functionality of the PDA coating, a permeation study was performed with gelatin microparticles coated for either 0, 2, 4, or 24 hours. The capsules were placed in a PBS solution with FITC labelled dextrans (250 kDa) for 24 hours and subsequently imaged using confocal microscopy to measure capsule fluorescence at 20 and 37 °C. There was a noticeable decrease in capsule fluorescence as coating time increases for the 20 °C condition, though minimal (**Figure S3A-B**). When assessing the 37 °C conditions, it became clear that stable coatings did not occur until after 4 hours of coating (**Figure 2C-D**). Even after 4 hours, faint silhouettes of capsules can be seen with a significant difference in fluorescence when compared to the room temperature condition (P < 0.0001), suggesting a leaky coating. However, after coating for 24 hours, there is a slight difference in permeability between the two temperature conditions. Therefore, 24-hour coatings were subsequently used for the rest of the work.

We next sought to verify that our capsules can encapsulate cargo. To measure encapsulation efficiency, loaded capsules (2 mg/mL of 10, 40, or 250 kDa FITC-Dextran) were synthesized identically but without any dopamine added to the coating solution. The particles were then left to shake for 4 hours without dopamine for only four hours to simulate the time until a stable coating would form and prevent further leakage (**Figure 2D**). Confocal microscopy images of capsules loaded with 250 kDa dextrans were first used to confirm loading (**Figure 2E**). The uncoated capsules were then collected, washed, and melted. A fluorescent spectrophotometer was used to measure the amount of FITC-dextran (**Figure 6.8A**) with a emission maximum of 517 nm. To quantify the dextran concentration, standard curves of FITC-dextrans at varied concentrations were made for each size (10, 40, and 250 kDa) (**Figure S4A-B**). Confocal microscopy images of capsules loaded with 250 kDa dextrans were first used to confirm loading (**Figure 2E**). The encapsulation efficiencies for 10, 40 and 250 kDa dextrans were 32 ± 2%, 32 ± 4%, and 30 ± 0.5% respectively (**Figure 2E**). Material loss most likely occurs during the coating process before a sufficiently thick enough coating is formed.

To measure the long-term stability of the coatings, FITC labelled bovine serum albumin (BSA) molecules were loaded into capsules and stored at 4 °C. These capsules were subsequently imaged under a confocal microscope over 28 days (**Figure S4C**). To combat the gradual photobleaching of FITC, the internal particle fluorescence was normalized to a solution of free FITC-BSA stored under the same conditions. A 94 ± 1% retention in fluorescence is seen over the 28 days (**Figures 2F**). To test the maximal loading capacity of our capsules, 10 wt% BSA (with 0.1% being FITC-BSA) was added to the gelatin precursor solution. A similar procedure to the encapsulation efficiency was performed on supernatants of dissolved non-coated capsules after 4 hours (**Figure S5A**). A standard curve was made for the FITC-BSA (**Figure S5B**) which gave a final BSA concentration of 39 ± 2 mg/mL of BSA (**Figure 2G**). To our knowledge, this is the highest loading of bioactive molecules in a microcapsule to date.

Microscale hydrogels have been widely used as delivery vehicles, scaffolds and support matrices for biofabrication.^22,23^ We next aimed to determine if a jammed suspension of capsules would behave similarly to standard microgel suspensions. To induce jamming, capsules were placed on 25 μm nylon filters to drain away excess water. Parallel plate rheology on jammed microcapsules revealed similar shear thinning behavior (**Figure 2H**) with a yield stress value of 40 Pa (**Figure S6A**). To demonstrate reversibility of their jammed state, 2-minute intervals of high (200%) and low (0.2%) strain were applied cyclically. Here, the jammed capsules can be seen to recover full mechanical stiffness near instantaneously for multiple continuous cycles (**Figure S6B**). To assess injectability, jammed microgels were loaded into 1 mL syringes to be extruded. Notably, when the particles have not drained enough water, the ink does not print as a single jammed line (Supplemental videos 1-3). If too wet, the ink forms small droplets at the nozzle tip. As the suspension loses more water, the droplet begins to shrink in size and small fragments of jammed particles can be printed. However once adequately dry and jammed, the suspension prints as a single unbroken line (**Figure 2I**). These findings correspond with those found in recent work by Alge and colleagues where unjammed microgel inks will expel water print poorly until a critical microgel concentration is reached.^24^ Finally, a fidelity assessment of 3D printed lines demonstrates high control of line with widths varying from 300 – 800 μm, with average capsule line width diameter less than 5% from of the nozzle’s inner diameter (**Figure 2J-K**). These studies demonstrate how the microcapsules can be used in multiple formats from single particles to high density suspensions for injection.

### 3.2 Glucose oxidase loaded capsules reduce dissolved oxygen levels

We next evaluated the ability to load functional enzyme into our new microcapsule system. First, glucose oxidase was loaded into microgel capsules (1 U/mL). A glucose oxidase activity kit with a fluorescent reporter was used to measure the hydrogen peroxide production activity of enzyme loaded microcapsules. Capsules (20 mg/mL) were added to wells in a plate reader and the fluorescent output (515 nm) was measured every 3 minutes for 45 minutes. When measuring the activity over time, the oxygen levels above the capsules reached a plateau after 3-6 minutes (**Figure 3A**). This is likely due to initial diffusion constraints as glucose substrates and peroxides need to freely diffuse into and out of capsules. After 45 minutes, capsules loaded with 1 U/mL of glucose oxidase had an activity of 46 mU/mL, providing a functional encapsulation efficiency of 5%. This discrepancy between these values and the 30% found for loaded BSA may be due to loss of functionality of some of the encapsulated enzymes during encapsulation and coating at elevated pH (~ 8.5). The substrates and products of the reaction also need to diffuse through the capsule wall measurement which can limit the reaction rate and the perceived activity. In addition, immobilized enzyme activity can decrease by up to 80% when compared to free enzyme.^25^ This underscores the importance of an encapsulation system with high cargo loading capacity.

**Figure 3.**
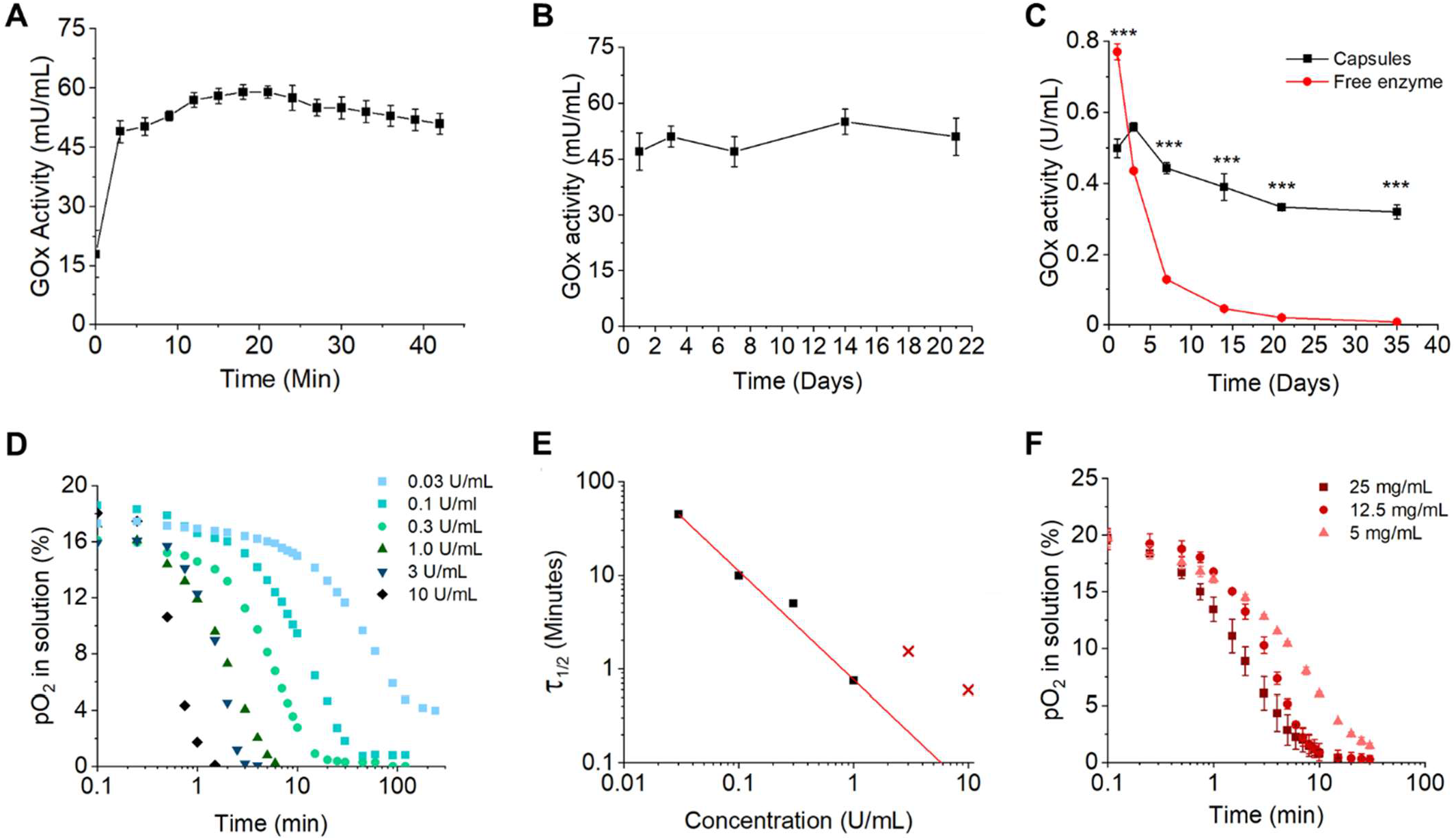
Characterization glucose oxidase activity in capsules. A) Glucose oxidase (GOx) activity overtime for glucose oxidase loaded microcapsules (1 U/mL) measured with a glucose oxidase activity kit on plate reader with 517 nm emission. B) Long term activity of glucose oxidase capsules stored at 4 °C in between measurements. C) Long term activity of glucose oxidase capsules and free enzyme stored at 37 °C in between measurements. D) Plot of the dissolved oxygen partial pressures for varied concentrations of glucose oxidase (with a 1:60 ratio with catalase) in cell media at room temperature. E) plot of the saturation half time versus glucose oxidase concentration. The two x’s correspond to data points that were not used for the linear fit. F) A plot of the dissolved oxygen partial pressures for capsules at varied concentrations (5, 12.5, or 25 mg/mL of glucose oxidase loaded capsules (1000 U/mL, 60,000 U/mL catalase)). ***P < 0.001, ANOVA. Error bars represent s.d.

Glucose oxidase is known for having high shelf life under proper storage conditions.^26,27^ To measure the shelf life of the enzyme loaded capsules at 4 °C in PBS, a batch of capsules had activity measured at 1, 3, 7, 14, and 21 days after synthesis. No noticeable difference in glucose oxidase activity was found after 21 days in storage demonstrating high stability. Next, glucose oxidase loaded capsules were stored in cell culture media (DMEM) at 37 °C for five weeks, along with a solution of free enzyme as control. After 1 week, the capsules lost only 11 ± 0.3% enzyme activity (P = 0.0094), whereas the free enzyme had a reduction of activity of 84.4 ± 0.1% (P < 0.0001). After 5 weeks, the capsules lost only 34 ± 0.4% enzyme activity (P < 0.0001), whereas the free enzyme had lost 99 ± 0.2% enzyme activity (P < 0.0001). Together, these results demonstrate how our microencapsulation system yields excellent enzyme stability providing a tool that could maintain hypoxic cell cultures for weeks.

With confirmation that capsules loaded with glucose oxidase can generate hydrogen peroxide, we next aimed to verify that encapsulated glucose oxidase, along with catalase, can reduce oxygen levels. For all tests, catalase was loaded at a functional activity of 60 times that of glucose oxidase. This ratio is to ensure that hydrogen peroxide produced by glucose oxidase is immediately extinguished by the catalase. This ratio is also consistent with previous work using enzymes to create hypoxic cell culture environments.^9,14^ To best simulate the oxygen level conditions for cell culture conditions, a dissolved oxygen (DO) probe was placed 7 mm deep (the depth of our hydrogels in a 24 well plate with 1 mL of liquid) into a 12 well plate loaded with 4 mL of cell culture media (**Figure S1**). A 6 mm stir bar was added to the well and rotated at 250 rpm to ensure enough fluid flow at the probe interface for accurate measurement while preventing disruption of the air-liquid interface. All measurements were performed at 21 °C. When testing at room temperature and pressure, the probe baseline read 9.07 mg/L of oxygen which, when using Henry’s law, gives a partial pressure of 21.0% oxygen.

The dissolved oxygen levels were measured for glucose oxidase concentrations of 0.03, 0.1, 0.3, 1, 3, and 10 U/mL. As expected, the dissolved oxygen levels were found to decrease as a function of the glucose oxidase concentration (**Figure 3D**). At each concentration, the oxygen saturation versus time relationship follows a sigmoidal curve shape. As the added enzymes first begin removing oxygen, there is a slight delay before the probe can register the reduction. And once the oxygen levels begin reaching lower saturation levels (< 1%), the enzymes begin to compete with the oxygen diffusion from the air-liquid interface. This is most apparent when comparing the 10 U/mL sample vs the 0.03 U/mL. Since some solutions do not reach 0% saturation, a different metric is needed for comparison between concentrations. Here, we define the saturation half time. This half time is defined as the time required for the dissolved oxygen levels in solution to reach half (10.5% pO_2_) of atmospheric oxygen concentration (a pO_2_ of 21%). The half times for each solution was plotted against the concentration of glucose oxidase to create a standard curve. The half time-activity relationship followed a power law model with high accuracy (R-square = 0.995) for activities of 1 U/mL and below (**Figure 3E and S7C**). Notably, the 3 and 10 U/mL solutions did not decay fast enough due to the limitation in speed of our oxygen probe. Therefore, they were not used for the best fit equation.

To measure the effect of catalase on the solution, pure glucose oxidase (0.3 U/mL activity) without catalase was added to the microcapsules. Here, the sample without glucose oxidase shifts the activity curve to the left, suggesting faster oxygen depletion. Calculation of the saturation half time gives 5.8 and 3.5 minutes, a 40% reduction, for the catalase and catalase free conditions respectively (**Figure S7A-B**). This fits with our expectations as catalase should lead to a reduction of oxygen depletion by 50%. Glucose oxidase and catalase loaded microcapsules of varied activities (1000 U/mL of glucose oxidase and 60,000 U/mL of catalase; 5, 12.5, and 25 mg/mL of capsules) were subsequently added to the same DO probe setup to measure the effect of encapsulation on activity. These capsules all reached hypoxic levels of oxygen tension (< 5% pO_2_ oxygen) by 15 minutes, with a half saturation time of 1.9 ± 0.4, 3.0 ± 0.3, and 5.9 ± 0.6 minutes for the 5, 12.5, and 25 mg/mL conditions respectively (**Figure 3F**).

Finally, a capsule loaded GelMa microgel suspensions (2 vol% capsules in the gel, 0.4mL of hydrogel volume; final of 2 mg/mL of capsules) was analyzed with the identical setup. These loaded microgel suspensions had a half saturation time of 23 minutes, reaching a plateau of 2% oxygen saturation after 2 hours (**Figure S7D**). These data suggest that hypoxic conditions can readily be reached with a low total volume fraction of capsules. One caveat with this methodology is that it can only read the oxygen levels in solution. It is well appreciated that both *in vitro* hydrogels and *in vivo* tissues can lead to reduced oxygen levels when thick enough.^28^ And even slight differences in depth of cell culture medium in flasks and wells can affect the oxygen levels to adherent cells.^6,29^ Therefore, the measurements of liquid at the surface likely underestimate the levels of hypoxia that cells experience within the microgel suspensions. Nonetheless, these measurements provide insight into the upper bound of glucose oxidase needed to establish these conditions.

### 3.3 Low oxygen environments perturb MSC function and tumor growth

With functioning oxygen modulating microcapsules, we investigated the effect of hypoxia on cell viability. Live dead assays were performed on days 1 and 5 for 3 cell lines: A375-P, Adipose derived stromal cells (ADSC), and human umbilical vascular endothelial cells (HUVECs). Capsules loaded with 1000 U/mL of glucose oxidase and 60,000 U/mL of catalase were added to GelMa microgel suspensions, used previously,^13^ (2 vol% of capsules) along with one of the cell lines (**Figure S8**). Here, 1000 U/mL inside the capsules would give 2 U/mL in total solution in the well of a 24 well plate. With a 5% enzyme functionality, this provides a rough activity of 100 mU of glucose oxidase activity in one well (1 mL of volume). Since all following *in vitro* studies were performed in 24 well plates with 1 mL of media, all future references of U/mL of enzyme will refer to the initial loading within the capsules for consistency.

After 1 day, there was a significant reduction in viability for the melanoma cell line, from 97 ± 1% to 65 ± 20% when hypoxic capsules were added, while the ADSCs and HUVECs had little difference (**Figure S8A-B**). By 5 days, nearly all A375-P cells were dead (16 ± 12%, P = 0.000017), while HUVEC cells had a drop in viability to 36 ±20 % (P = 0.0016) (**Figure S8C**). Surprisingly, the ADSCs showed no decrease in viability even after 5 days (93.7 ± 0.3 vs 92.5 ± 5.5%, P = 0.999). However, the ADSCs, along with the HUVECs, showed spreading in the control condition, but no noticeable cell spreading in the glucose oxidase condition (**Figure 8SA**). Data from the dissolve oxygen probe showed these capsules exhibit an activity around 0.3 U/mL which gives a < 1% oxygen saturation after 20 minutes at the surface of the gel. Thus, while some cells may survive at this level of hypoxia, cell function is significantly impaired.

To explore the effect of hypoxia on intracellular HIF1-alpha, microgel suspensions were loaded with ADSCs (1 × 10^6^ cells per mL) with either no capsules, or capsules with 100, 250, or 1000 U/mL of glucose oxidase (2 vol%; 6,000, 24,000, or 60,000 U/mL of catalase respectively). At both days 1 and 3, significant expression of HIF1-alpha can be seen in in the capsule conditions but not the control (**Figure 4A**). Here, the control sample had roughly 5% of cells with expression while the 1000U/mL glucose had 73% (P = 0.0013) (**Figure 4B**). Hypoxia has been shown to have an effect on the extracellular vesicle (EV) output of mesenchymal stem cells.^30,31^ To measure if our capsule induced hypoxia had any effect on ADSC extracellular vesicle output, ADSCs were embedded into 3D microgel suspensions with capsules containing 0, 30, or 250 U/mL of glucose oxidase (1 × 10^6^ cells per mL, 2 vol% capsules in hydrogel). After 72 hours in serum free media, the supernatants were collected, and the EVs were isolated and analyzed. No differences were found in the concentration of exosomes across all three conditions (**Figure 4C**). However, there was a significant increase in mean vesicle diameter for the hypoxic condition compared to the control (P = 0.010) (**Figure 4D-E**). This correlates well with previously reported work.^32^ The presence of CD9, a known surface marker for exosomes, in the EVs was also confirmed via western blots (**Figure S9**). Overall, this data suggests that the ADSCs are indeed experiencing and responding to the low oxygen environment, leading to elevated secretion of extracellular vesicles.

**Figure 4.**
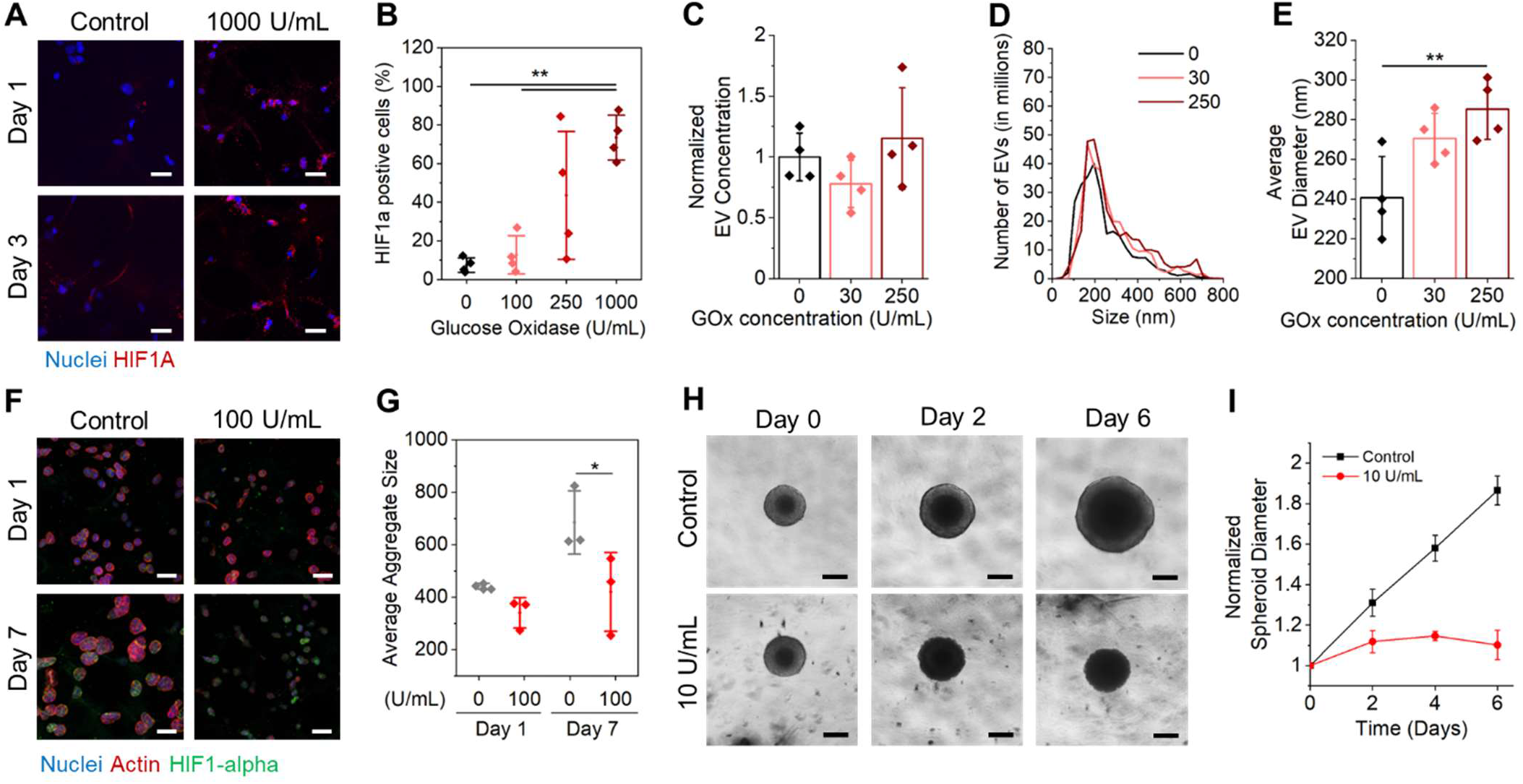
Effect of Hypoxia on ADSCs and tumor cells. A) Confocal z-stack projections of ADSC cells (1 × 10^6^ cells per mL) stained with Hoechst (Nuclei, blue), phalloidin (Actin, not shown), and HIF1-alpha (red) after 1 or 3 days in culture in microgel suspensions (10 wt%, 1% filler) with or without 2 vol% of glucose oxidase capsules (1000 U/mL, 60,000 U/mL of catalase). B) Cell counts per replicate region of HIF1-alpha positive cells for each of the glucose oxidase concentrations (capsules loaded with 0, 100, 250, and 1000 U/mL). Concentrations (C), size plots (D), and average diameters (E) of EVs isolated from ADSCs in microgel suspensions with capsules loaded with varied glucose oxidase concentrations. F) Confocal z-stack projections of A375-P cells stained with Hoechst (Nuclei, blue), phalloidin (Actin, red), and HIF1-alpha (green) after 1 or 7 days in culture in microgel suspensions (Medium, 10 wt%, 1% filler) with or without 2 vol% of glucose oxidase capsules (100 U/mL, 6,000 U/mL of catalase). G) Plots of average A375-P aggregate size per replicate overtime. H) Optical images of MCF7 breast cancer spheroids embedded in Geltrex over 6 days with no capsules or with capsules loaded with an equivalent of 10 U/mL. I) Growth curves of breast cancer spheroids diameters. Scale bars: 50 μm (A, F), 200 μm (H). *P < 0.05, **P < 0.01, ANOVA. Error bars represent s.d.

When sufficiently large enough, all cancerous tumors exhibit hypoxic cores.^33^ Aberrant tumor paracrine signaling networks disrupt nearby vasculature, leading to leaky vessels and poor nutrient and oxygen delivery. This disruption can generate hypoxic cores with a tumor volume as they continue to grow. These hypoxic cores have been implicated in tumor progression, including higher cancer survival, aggressiveness, and more frequent incidence of metastasis.^33–35^ Therefore, model systems that can replicate hypoxic microenvironments in the laboratory are important research tools for fundamental research and drug development.

To measure the effect of hypoxia on melanoma cells, we embedded A375-P human melanoma cells in GelMa microgel suspensions (between particles) with a filler of 1 wt% GelMa in the microgel interstitial space. These suspensions were subsequently loaded with microgel capsules that contained 0 or 100 U/mL of glucose oxidase (and 6,000 U/mL of catalase), 10% of the enzyme activity that led to total cell death of the A375-Ps in the live dead study. In the control condition, A375-P cells can be seen to grow in clusters that expand in size by 55% (P = 0.07) over the course of 7 days (**Figure 4F**). In contrast, the cell clusters in microgel suspensions with hypoxia inducing capsules increased in size by only 23% (P = 0.77) and 61 ± 20% the size of the control condition (P = 0.049) (**Figure 4G**). Notably, this model used single cells which are less reminiscent of *in vivo* tumors. Therefore, we next explored tumor spheroids to evaluate if hypoxia would affect tumor outgrowth. Here, MCF-7 breast cancer spheroids were embedded into Geltrex with 0, 10, 30, or 100 U/mL capsules loaded. For the conditions with 30 or 100 U/mL capsules, the spheroids did not survive prolonged culture. For the 10 U/mL condition, there was slight growth through the first two days, after which the spheroids died (**Figure 4H-I**). These findings were replicated when using a more aggressive breast cancer cell line, MDA-MB-231, where there was little tumor growth for the hypoxic condition over 6 days (**Figure S10**).

Together, our findings suggest that tumor cells within our in *vitro* models are sensitive to hypoxia, eventually leading to cell death. This increased susceptibility of tumor cells to apoptosis under hypoxia in vitro has previously been demonstrated, with the level of susceptibility varying from cell line to cell line.^36,37^ Yao and coworkers speculated that the level of p53 competence of the cancer cells influences their sensitivity to hypoxic stress.^36^ Nonetheless, our results suggest that it may be difficult to draw direct conclusions of *in vivo* tumor response using *in vitro* hypoxia models.

### 3.4 Hypoxic microcapsules induce angiogenesis *in vitro* and *in vivo*

It is well understood that oxygen levels play a role in endothelial cell signaling pathways.^38^ Under low oxygen tension, endothelial cells are driven to undergo angiogenesis to create more vessel structures to supply more oxygen. This has also been previously demonstrated using hydrogels with oxygen reducing enzymes tethered to the hydrogel backbone.^11^ Here we aimed to measure the effect of hypoxia on vascular endothelial cells. First, HUVECs were seeded in GelMa microgel suspensions (0% Filler, 5 × 10^6^ cells per mL) with varied concentrations of glucose oxidase loaded capsules and cultured for 24 hours before snap freezing the gels and collecting cellular mRNA. RT-qPCR analysis showed an upregulation in VEGFa levels with increasing levels of hypoxia, with a 2-fold difference for the 250 U/mL condition compared to the control condition (**Figure 5A**) (P < 0.0001). There was a simultaneous decrease in ANGPT-1 by 10-fold (**Figure 5B**) and no statistical difference in MMP1 gene expression (**Figure S10A**). Previous work has demonstrated that hypoxia induces an upregulation in VEGFa and a downregulation in ANGPT-1 in human endometrial cells.^39^ This is in contrast to other literature showing an upregulation of ANGPT-1 in vascular cells,^11^ and pericytes under hypoxic conditions.^40^ Park et al. have shown that this upregulation can be acute and transient.^41^ The role of ANGPT-1 is primarily vascular protective, preventing endothelial cell death and preserving the structural integrity of vasculature.^42^ It is possible that in the first 24 hours the dispersed cells increase VEGFA expression due to hypoxia, but without the paracrine mechanism that is vascular protective. To test whether our endothelial cells would still undergo angiogenesis in this system, the previous experiment was repeated but allowed to proceed for 5 days. After 5 days, HUVECs in enzyme capsule loaded conditions (100 and 250 U/mL of glucose oxidase) began to form interconnected tubes (**Figure 5C and S12A**). For quantification, the angiogenesis analyzer tool in ImageJ was used (**Figure S12B**). When comparing the two conditions, the 100 and 250 U/mL conditions had 430% and 470% increased master segment length (0 – 100: P = 0.019; 0 – 250: P = 0.0070) and 320% and 460% more master segments respectively (0 – 100: P = 0.227; 0 – 250: P = 0.022). However, when measuring average branch length (**Figure S11B**), as well as mesh area (**Figure S11C**), there was a trend suggesting that the 100 U/mL had thicker and healthier vessels, which matches the qualitative observations from the confocal images (**Figure S12A**).

**Figure 5.**
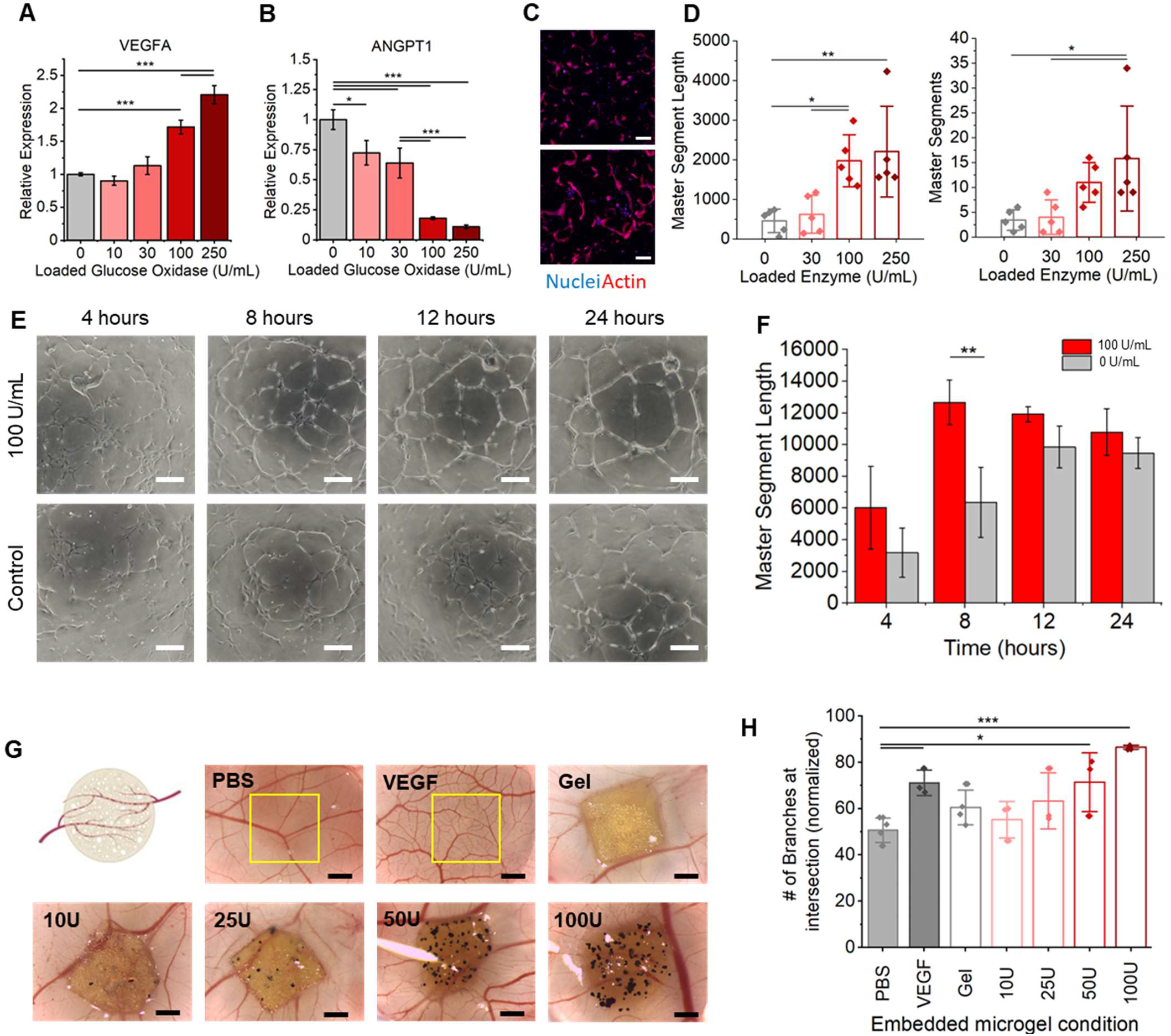
Effect of hypoxic capsules on endothelial cells. Plots of relative expression of VEGFA (A) and ANGPT-1 (B) for HUVECs in microgel suspensions with varied amounts of hypoxic capsules for 24 hours. Plots normalized to GAPDH. C) Confocal z-stack projections of HUVEC (5 × 10^6^ cells per mL) cells stained with Hoechst (Nuclei, blue), and phalloidin (Actin, red), after 5 days in culture in microgel suspensions (10 wt%, 0.5% filler) with or without 2 vol% of glucose oxidase capsules (100 U/mL, 6,000 U/mL of catalase). D) Plots of master segment length (right) and number (left) and lengths for control and glucose oxidase capsule conditions calculated using the angiogenesis analyzer plugin for ImageJ. E) Bright field images of HUVEC tubulogenesis over the course of 24 hours. F) Analysis of master segment length from the tubulogenesis assay. G) Dissecting microscope images of representative CAM assay egg embryos after 4 days of incubation with or without microgel suspensions loaded with glucose oxidase capsules. H) Quantification of the number interacting branching points from the CAM assay. Scale bars: 50 μm (C), 100 μm (E), 500 μm. (G). *P < 0.05, **P < 0.01, ***P < 0.001, ANOVA. Error bars represent s.d.

To further confirm these angiogenesis results, a tubulogenesis on Matrigel assay was performed using the 0 U/mL and 100 U/mL capsules conditions (**Figure 5E**). These results demonstrated a temporal response to hypoxia, where the hypoxic condition reached its maximum master segment length (**Figure 5F**), number of master segments (**Figure S11D**), and number of master junctions (**Figure S11E**) 4 hours earlier than the control conditions. These differences abated by 24 hours as is expected with endothelial tubulogenesis experiments on Matrigel. As a final confirmation of the hypoxia *in vivo*, we ran a Chick Chorioallantoic Membrane (CAM) assay, a well-established *in vivo* angiogenesis assay. Here, microgel suspensions loaded with varied concentrations of 100 U/mL capsules were placed onto the CAM of developing chicken embryos. After 4 days of sample exposure, the number of branches intersecting the edge of the gels was analyzed. No statistical differences were found between the PBS and the gels loaded with 0, 10, or 25 U/mL enzyme capsules. However, there was a significant difference between the PBS and the 50 and 100 U/mL conditions (PBS – 50: P = 0.02; PBS – 100: P = 0.0002). Further, the 100 U/mL condition enhanced angiogenesis with 43% and 22% more intersecting vessels relative to the 0 U/mL gels and the VEGF positive control, respectively. Overall, these data suggest that hypoxia induced upregulation of VEGFa leads to angiogenesis in 2D, 3D, and *in ovo* models. The ability to fine tune the concentration of capsules and enzyme allows for a robust way to investigate subtle changes in the levels of oxygen tension and how this perturbs angiogenesis and associated processes.

### 3.5 Ceria loaded capsules to reduce cellular oxidative stress

Finally, to demonstrate cargo flexibility we evaluated the loading of gas modulating nanozymes within the capsules. We chose cerium oxide nanorods (Nanoceria) as our candidate payload since these nanozymes have found widespread use as *in vivo* therapeutics by attenuating reactive oxygen species.^43^ Nanoceria has catalase (**Figure 6A**), ATPase, and peroxidase activities depending on its valence state and buffer pH,^44,45^ and it can reduce the oxidative stress on cells in vitro.^46^ Nanoceria’s enzymatic mimicry make it an attractive cargo as enzymatic substrates could diffuse in and out of the capsule wall while the nanoceria remain entrapped.

**Figure 6.**
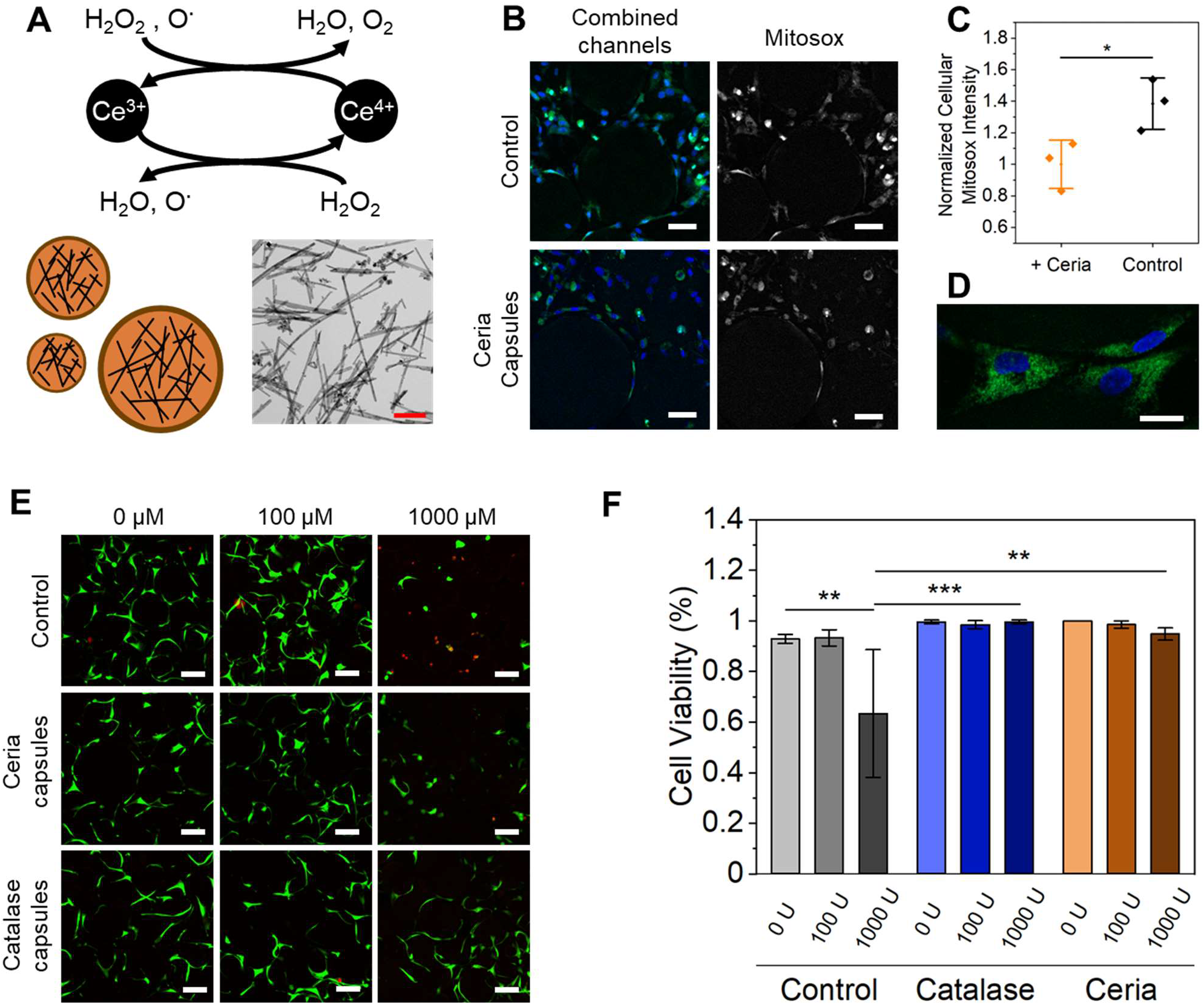
Nanoceria rods loaded into microcapsules. A) Reaction scheme of nanoceria’s catalase activity (Top), schematic of nanoceria in capsules (left), and TEM image of nanoceria rods (Right). B) Confocal z-stack projection of live ADSCs stained with Hoechst (Nuclei, blue) and MitoSOX (superoxide species, green) in either control microgel suspensions (Medium, 10 wt%, 1% filler) or in suspensions with 2 vol% ceria loaded microcapsules (5 mg/mL). B) Normalize intensity plots of cellular MitoSOX levels take from the fluorescence images. D) High resolution image of MitoSOX and Hoechst stained ADSCs. E) Confocal microscopy z-stack projections of live dead stained ADSCs (1 × 10^6^ cells per mL) after 24 hours of exposure to H_2_O_2_ (0, 100, 1000 μm) in microgel suspensions (Medium, 10 wt%, 0% filler) with either no capsules, 2 vol% ceria loaded capsules (5 mg/mL), or 2 vol% catalase loaded capsules (60,000 U/mL). F) Cell viability plots of the data captured on confocal. Scale bars: 200 nm (A), 5 μm (D), and 50 μm (B, E). *P < 0.05, **P < 0.01, ***P < 0.0001, ANOVA. Error bars represent s.d.

Nanoceria was synthesized using published methods to produce nanorods with lengths of 50 nm (**Figure 6A**). To quickly verify the functionality of this nanoceria, the particles were suspended in a McIlvaine buffer solution (0.1M) at varied pH (3, 5, and 7). A solution of an organic dye that changes color when oxidized (2,2’-azino-bis(3-ethylbenzothiazoline-6-sulphonic acid) (ABTS)) by cerium oxide was paired to measure the peroxidase mimicking activity of the nanoceria in solution.^47,48^ To quantify the pH and nanoceria activity relation, ceria and ABTS were added to an array of buffers with varied pH (4, 5, 6, 7, and 8, 2mM ABTS, 0.5 mg/mL ceria) and were left to react overnight. The absorbance spectra showed an order of magnitude higher activity of nanoceria at a pH of 4 versus 6, with a plateau of activity beyond pH 6 (**Figure S13B**). This correlates well with the pH response found in the literature, demonstrating that our nanoceria behaves as expected.^47^

To verify that nanoceria can be loaded into capsules, we conjugated fluorophores to the ceria particles for visualization. Successful silanization of nanoceria was verified using FTIR where appearance of the 1050-1100 peak for the silanized samples represents the Si-O-Si bond stretching (**Figure S14A**).^49^ After particle resuspension in ethanol, a solution of FITC (1 mg/mL) was added for direct conjugation to the terminal amine groups. Confocal microscopy images demonstrate successful fluorophore conjugation as well as successful loading of nanoceria into microgel capsules (**Figure S14B**). Typically, nanoceria loaded in high concentrations above 2-5 μg/mL would be cytotoxic to cells.^50^ However, by adding the nanoceria to capsules, we can reach high concentrations that maintain adequate functionality while protecting cell viability. To assess the effect of a high concentration (5 mg/mL) of nanoceria on cell viability, a live dead assay was performed with three cell types: A375, ADSC, and HUVEC (**Figure S15A**) Nanoceria loaded microgel capsules were added to GelMa microgel suspensions at 2% of the final volume.

Live dead staining at both days 1 and 5 indicate a viability above 90% for all cells lines with ceria capsules (**Figure S15B-C**). One functional use of nanoceria in cell culture is the reduction of superoxide species (O_2_^•-^).^51^ Cellular superoxides are typically formed in mitochondria during cellular respiration.^52^ If they accumulate too much, they can generate high levels of hydrogen peroxide and hydroxyl radials which disrupt cell activity and led to cell apoptosis.^53,54^ To combat this damage, cells use the enzyme superoxide dismutase (SOD). ^55^ In humans, up to 99% of SOD is found in the extracellular matrix and blood plasma.^55^ Nanoceria has also been shown to reduce superoxide tension in cells in culture and improve cell viabilities.^46^ However, studies have only demonstrated super oxide protection when nanoceria particles are endocytosed into cells. Therefore, we hypothesized that high concentrations of nanoceria loaded within microcapsules, simulating extracellular SOD enzymes, could also reduce the superoxides within the surrounding matrix. First, ADSCs were seeded in either capsule free microgel suspensions or microgel suspension with ceria-loaded capsules (5 mg/mL of nanoceria, 2 vol% capsules). After 5 days, the cells were stained with Hoechst and MitoSOX, a live cell dye that penetrates cell membranes and fluoresces when exposed to superoxide species (**Figure 6B**). Analysis of confocal z-stacks showed a 37% reduction in cellular MitoSOX intensity in cultures with ceria-loaded capsules (P = 0.040) (**Figure 6C**). To confirm correct staining, high resolution images were taken that show a punctate speckled appearance (**Figure 6D**), matching with mitochondrial stains.^56^

Ceria nanoparticles also have potential therapeutic use during cancer treatment. A common cancer therapy is radiation where a focused blast of a photon or particle beam is aimed at the tumor site.^57^ However, while these blasts are typically well targeted, they can affect the surrounding matrix, leading to unwanted damage of healthy tissues, including the generation of hydrogen peroxides.^58^ This large influx of peroxides can cause significant damage to surrounding tissues via radical hydroxide formation and subsequent DNA damage. Previous work has created in vitro models of this peroxide generation to assess the cytoprotective effect.^59^ Inspired by these reports, we aimed to measure the cytoprotective effect of nanoceria against peroxides. Here, ADSCs were embedded in microgel suspensions and subjected to 24 hours of culture with varied concentrations of hydrogen peroxide (0, 100, and 1000 μM). For each concentration range, replicates were made with ceria capsules loaded into the suspension (5 mg/mL) at 2% volume fraction. As a negative control, no ceria capsules were added to the suspension; as a positive control, a high concentration of catalase enzyme (1000U/mL) was loaded into microgel capsules and added to the microgel suspension (**Figure 6E**). When quantifying cell viability, all conditions with equal to or less than 100 μM of H_2_O_2_ had cell viabilities above 90% (**Figure 6F**). When exposed to 1000 μM, the negative control samples had 63% live cells compared to 99.5% (P = 0.00047) and 95% (P = 0.0027) for the catalase and ceria conditions respectively. In addition, the ceria capsule conditions appeared to have significantly less cell spreading than their catalase counterparts. This data suggests that the ceria nanoparticles are able to remove hydrogen peroxides from solution and protect cell viability, but potentially at a slower rate than catalase, leading to lower cell spreading and slower recovery.

## 4. Conclusions

This work presents a novel way to spatially control hypoxic conditions in any 2D or 3D hydrogel system. The microcapsule system enables high loading of enzymes with prolonged shelf life and maintenance of activity, which allows for control of hypoxia in cell culture for several weeks. Tuning the capsule concentration or initial enzyme concentration allows for fine control over the oxygen levels across a 3D cell culture, and across different samples in a well plate, towards integration in high throughput assays. Hypoxic conditions were shown to attenuate MSC EV output, as well as dimmish tumor growth characteristics. Hypoxic capsules also promoted an upregulation in endothelial VEGFa which led to enhanced angiogenesis in 2D, 3D, and *in ovo* models. This approach can also be adapted to nanozymes, other enzyme systems, or gas releasing hydrogels for use in a broad range of applications. The injectability of the capsules suggests potential use for direct *in vivo* therapy or as a cytoprotective agent to attenuate damage due to radiation therapy. The flexibility of the microcapsule format provides scope for use as a reservoir for inclusion in topical treatments and gel-based bandages, where gas attenuation may have clinical utility (e.g., wound healing). In addition to these practical applications, we anticipate this system will provide a basis for future work investigating the role of oxygen deprivation on cell signaling pathways and functional cellular outputs.

## Acknowledgements

This work was supported through funding from the National Health and Medical Research Council Grant # APP1185021, the Australian Research Council Grant # DP210103654, and the National Cancer Institute of the National Institutes of Health Grant # R01CA251443. We acknowledge the help and support of staff at the Katharina Gaus Light Microscopy Facility and the Biological Specimen Preparation Laboratory of the UNSW Mark Wainwright Analytical Centre.

## Conflicts of interest

There are no conflicts of interest to declare.

## Notes

### Competing Interest Statement

The authors have declared no competing interest.

