## Supplemental for "Gas modulating microcapsules for spatiotemporal control of hypoxia"

### Supplementary Methods

##### Gelatin methacryloyl (GelMa) synthesis

GelMa was synthesized as previously described.[60] Briefly, Type A gelatin from porcine skin (Bloom strength 300) was dissolved at 10% (w/v) in 1X phosphate buffered saline (1X PBS, pH 7.4) under stirring at 50°C. 5% (v/w) methacrylic anhydride was added and the mixture stirred for 90 minutes. The solution was diluted two-fold with 1X PBS and centrifuged (3000 g, 3 minutes) to remove unreacted methacrylic anhydride particulates. Following this, it was transferred into 14 kDa cutoff cellulose dialysis tubes and dialyzed at 40 °C for 5-7 days against deionized water. The dialyzed solution was lyophilized for 5-7 days, and the resulting powder was stored at -20 °C.

##### Scanning electron microscopy (SEM)

Prior to imaging, microcapsules were air dried at 40°C until no water remained. The samples were subsequently fixed to aluminum stubs with adhesive carbon discs. The samples and stubs were then sputter coated once with platinum (Emitech K575x Pt sputter coater). A Hitachi S3400 (7-10 kV, probe current of 40) scanning electron microscope was used for all imaging.

##### Particle permeation study

Four batches of microcapsules were formed with 0, 2, 4, and 24 hours of coating. Then, 25 mg of each type of microcapsule was added to a solution of 250 kDa Fluorescein isothiocyanate (FITC) dextran (100 µg/mL). The solutions were then stored at either room temperature or 37 °C for 18 hours and subsequently imaged on a confocal microscope (Nikon A1). The fluorescence level of 40-100 particles was then averaged and normalized to the background fluorescence from each image.

##### 3D printing Jammed Microcapsules

Coated particles were first filtered using a 180-micron metal sieve to remove any particles that had aggregated together into a clump. The particles were then placed onto a 25 µm nylon filter over paper towels to remove excess water. The microcapsules were then scoped up with a spatula and loaded into a 1 mL Livingston syringe for printing. The syringe was loaded into the Replistruder head of a modified Lulzbot mini 2 3D printer. Three nozzle sizes of Nordson EFD steel tip nozzles (23-gauge, 22-gauge, and 18-gauge) were used for measuring the fidelity of the 3D prints. Eight lines were printed from each needle gauge which were imaged under an optical microscope and analyzed using ImageJ. FITC-Dextran capsules (40 kDa Dextran, 10 µg/mL) were used for confocal imaging. For microgel suspension printing, an 18-gauge nozzle was used to print 1-5 µL of capsule droplets. All code used for 3D printing can be found at https://github.com/tmolley2/Gradient-Microgels.

##### Silanization and FITC conjugation of Ceria nanoparticles

Nanoceria particles were synthesized as described previously.[44] For salinization, 100 µg of nanoceria was added to 9 mL of 100% ethanol and sonicated for 10 minutes. Once particles were dispersed, 1 mL of (3-Aminopropyl) triethoxysilane (APTES) was added and the solution was placed on an orbital shaker at 300 rpm for 1.5 hours. The particles were centrifuged a 15000 g for 10 minutes and washed with 100% ethanol. This was repeated twice more. The functional groups on the bare and silanized ceria nanoparticles were determined by Fourier transform infrared (FTIR) spectroscopy (PerkinElmer FT-IR) in transmission mode (16 scans, wave-number range 4000–450 cm−1).

For FTIC conjugation, 1 mg/mL of FTIC was first dissolved in DMSO. While dissolving, 100 µg of silinated ceria nanoparticles were sonicated in 100% ethanol (8mL) to disperse. Next, 2 mL of the FITC/DMSO solution was added and the particles were wrapped in foil and placed on an orbital shaker at 300 rpm for 4 hours. The particles were centrifuged a 15000 g for 10 minutes and washed with 100% ethanol. This was repeated twice more before storing the particles at 4 °C in ethanol inside fresh centrifuge tube wrapped in foil.

#### Characterization of nanoceria peroxidase activity

Nanoceria was first centrifuged (15,000 g for 5 minutes) and air dried overnight in a fume cupboard. The ceria was weighed out into an Eppendorf tube and resuspended in DI water to 10 mg/mL. Mcilvaine buffers was prepared at a pH of 3, 4, 5, 6, 7, and 8 (0.1M). A solution of 2mM of 2,2'-azino-bis(3-ethylbenzothiazoline-6-sulfonic acid) (ABTS) was made in each buffer solution. For the qualitative cuvette test, nanoceria was subsequently added to each buffer at a final concentration of either 0.1 or 1 mg/mL. The solution was left to sit at room temperature for 24 hours prior to photographing. For the plate reader test, 95 µL of each buffered ABTS solution (pH of 4, 5, 6, 7, and 8) was added to seven wells in a 96 well plate. Then, 5 µL of ceria solution was added (final concentration of 0.5 mg/mL) to four of the wells, while three wells had 5 µL of the corresponding blank buffer added. The well plate was then left to sit overnight at room temperature before measuring their absorbance spectra at 415 nm on a plate reader (CLARIO Star Plus).

#### Cell viability with loaded microcapsules

Two microcapsule formulations were created, loaded with either 5 mg/mL of nanoceria or 10 mg/mL of glucose oxidase (1,000 U/mL) and 6 mg/mL of catalase (60,000 U/mL). Microcapsules were first washed four times with sterile 1X PBS containing 1% Penicillin/streptomycin. Three medium-sized microgel suspensions with 1 wt% filler, 10 wt% GelMa particles, and 0.05 wt% LAP were hydrated. One condition had no capsules, one had 20 mg/ml of ceria capsules, and one had 20 mg/mL of glucose oxidase capsules. Each suspension was split into three tubes and loaded with either ADSCs (1 × $10^6$ cells per mL), A375-P (3 × $10^6$ cells per mL), or HUVECS (5 × $10^6$ cells per mL). Next, 90 µL of suspension was added to 6×2.5×2.5 mm molds and photocrosslinked (with 395 nm UV light at 40 mW/cm$^2$) for 60 s. The gels were then cultured in 24 well plates with 1 mL of media for 1 or 5 days with media changes on day 1 and 3. For staining, the media was removed, and the sample were washed once with 1X PBS. A 1X PBS solution of Calcein AM (2 µM) and Ethidium Homodimer-1 (4 µM) was added (500 µL) to each well and the gels were incubated at 37 °C for 35 minutes. Next, the samples were washed with 1X PBS, placed back in the incubator for 5 minutes, and washed again before imaging on a Zeiss LSM 800 confocal microscope. To quantify cell viability, image analysis was performed with ImageJ. Briefly, each image was split into the live and dead channels, and each channel was thresholded to select only the cell volumes. The Analyze particle's function (0.3-1.0 sphericity, 40 µm$^2$ area) was then applied and the ratio of live to dead cells was used to determine the viability for each sample. The viabilities for three samples were then averaged.

#### Oxidative stress protection with nanoceria

Microcapsules were prepared with 5 mg/mL of ceria nanoparticles. Two microgel suspensions using medium-sized 10 wt% GelMa particles, and 0.5 wt% filler, were prepared with 0.05 wt% of LAP: one with 20 mg/mL of capsules, and one with no capsules. ADSCs were loaded into each suspension at 1 × $10^6$ cells per mL, and the suspensions were UV crosslinked in the standard plastic molds. The crosslinked suspensions were placed in 24 wells plates with 750 µL of media and subsequently cultured for five days with media changes on days 1 and 3. Then, 10 µg of MitoSOX was dissolved in 13 µL of DMSO before being diluted into 9 mL of 1X PBS. A 1mg/mL solution of Hoechst 33342 was diluted into the 1X PBS as well with a dilution ratio of 1:5000. Cell media was removed from the wells and replaced with 1 mL of the 1X PBS solution. The plate was placed back into the incubator at 37 °C for 40 minutes before the 1X PBS solution was removed and the samples were washed twice. The samples were then imaged on a confocal microscope (Zeiss LSM 800) where z-stacks of 100 µm (10x, 51 slices) were taken. Analysis was then performed in ImageJ. Here, a z-projection image of the stack was made, the MitoSOX channel was duplicated, and the copy had the background subtracted, (rolling ball radius of 20 µm). The image was then thresholded to capture the cell regions and a binary map was applied. These regions were converted into a selection and loaded into the ROI manager. The ROI selection was then applied to the original copy and the mean grey value of all the cells was calculated. This was repeated for each different stack of the three replicates for the control and microcapsule suspensions.

#### Dissolved oxygen concentrations

Dissolved oxygen measurements were taken using a Waterproof Dissolved Oxygen Meter (Hanna Instruments Pty Ltd, HI98193). Prior to use, the probe was turned on and allowed at least 15 minutes to condition to the room's temperature and pressure. Each day, a fresh 0 % oxygen solution was made (provided with the probe) and the probe was recalibrated after conditioning but before use. To simulate similar conditions used for cell culture experiments, the probe was place 7 mm deep into a 4 mL solution of DMEM high glucose media in a 12 well plate (**Figure S18**). A small 6 mm stir bar was also added and allowed to stir at 250 rpm to ensure enough fluid flow for accurate probe measurement readings. The probe was allowed to sit in the solution for 5 minutes to establish a baseline. Solutions of either pure glucose oxidase, or glucose oxidase and catalase, were made at a concentration of 10 mg/mL glucose oxidase (1000 U/mL) and 6 mg/mL of catalase (60,000 U/mL) in a 50mM sodium acetate buffer at a pH of 5.0. The solutions were then diluted to the concentrations needed as added to the stirring media solution. As soon as the solution was added, a time was started and measurements were recorded at recorded intervals for 120 minutes. If the solution reached 0 before then, then the recording was stopped. For the 0.03 U/ml glucose oxidase solution, 180 and 240 minutes were also recorded. A Saturation of 100% oxygen was defined as 9.07 mg/L given that it was the baseline value on the probe for our laboratory.

For capsule measurements, 50 mg of capsules, loaded with 10 mg/mL of glucose oxidase and 6 mg/mL of catalase, were added to the media solution and measurements were recorded. Two more replications were performed. For the capsule gel solution, a microgel suspension was made using medium-sized 10 wt% GelMa particles, 0.5 wt% filler, 0.05 wt% of LAP, and 20 mg/mL of the above glucose oxidase and catalase capsules. Four gels of 90 µL (identical to gels seeded with cells) were crosslinked and added to the 4 mL to a stirring media solution with the probe to simulate the same concentration of capsules to media used in cell experiments. Measurements were then taken with the same method as above.

To calculate the saturation half time value, the 20 to 60% saturation values were plotted, and a linear fit was used to calculate the 50% saturation value. The saturation half time values were then plotted on a log-log plot against glucose oxidase concentration, and a line of best fit was created using an allometric power law fitting.

#### Western Blotting

Identification of exosomes was confirmed by Western blot. Exosome lysates were prepared using RIPA buffer supplemented with a protease inhibitor (Thermo Fisher Scientific, Australia) and quantified by Pierce™ BCA Assay. Ten µg of exosome lysates were separated by SDS-PAGE, transferred onto a PVDF membrane, probed overnight at 4C with a primary antibody for CD9 (1:1,000 Cell Signaling Technology). and incubated with a secondary HRP-conjugated antibody (Abcam). Blots were visualised using Pierce™ ECL Western blotting detection reagents (Thermo Fisher Scientific, Australia), scanned using an ImageQuant LAS400 and quantified xxx Tom

#### Tumor spheroid growth models

For tumour spheroid formation, MCF-7 and MDA-MB-231 cells were seeded at 2000 and 4000 cells per well, respectively, in an ultra-low attachment round bottom 96 well microplate with a media change after 48h. At 72h post formation, the media was carefully removed, the spheroids were embedded in Geltrex loaded with 0, 10, 30, or 100 U/mL capsules (n=6 per condition) and grown under normal culture conditions for an additional 6 days. Cell culture media was changed every 48h and brightfield images were taken every 48h on an inverted light microscope (Olympus CKX53, 4x objective). Spheroid growth and outgrowth were quantified via ImageJ using the mean spheroid diameter at each time point.

#### Extracellular vesicle (EV) isolation and characterization

For EV isolation, cells were seeded in glucose oxidase capsule laden microgels and supplemented with serum-free media (Gibco StemPro™ MSC SFM, A1033201). EVs were isolated from cell-conditioned medium after 24hrs of culture using Total Exosome Isolation Reagent (Invitrogen, 4478359) according to the vendors instruction. In brief, cell conditioned media was centrifuged at 2,000 RCF for 30 minutes to remove cell debris and the supernatant removed. Total Exosome Isolation Reagent (500 μL) was added to the collected supernatant and incubated overnight. Samples were then centrifuged at 10,000 RCF at 4°C for 1 hour and the supernatant discarded. The obtained EV pellets were resuspended in 1 mL of particle free PBS.

EV size distributions and concentrations were determined by nanoparticle tracking analysis (NTA) using a NanoSight NS300 instrument (Malvern Technologies) that was configured using a high sensitivity CMOS camera and 488 nm laser. Samples were diluted (1:10) in particle free PBS and analyzed at 25 °C. Three 30 second videos were obtained using a camera level of 12. The obtained data was analyzed using NTA 3.2 software using a detection threshold of 7.

#### Nanoceria formulation and TEM characterization

Nanoceria was prepared using a hydrothermal synthesis. Briefly, equal amounts of 0.1M cerium nitrate and 10M sodium hydroxide solutions were combined under magnetic stirring at room temperature for 15 minutes. The solution was added to a steel autoclave and placed inside an oven at 120 ˚C for 24 hours, following which it was cooled to room temperature. The nanoparticles were separated and washed ten times with deionized water via centrifugation at 10 000 rpm for 10 minutes. The wet nanoparticles were stored in deionized water at room temperature until required for use. Nanoparticle morphology was characterized by Transmission Electron Microscopy (FEI Tecnai G2 20 TEM) at various magnifications after drop-casting a diluted nanoparticle solution onto a 3 mm copper TEM grid with a thin carbon film.

#### CAM assay

Fertilized black chicken eggs were supplied by Australian SPF services, VIC, Australia. All procedures were conducted in accordance with animal ethics protocols approved by the UNSW animal ethics committee (21/18B). All experiments were performed under aseptic conditions. On embryonic development day 0 (E0), eggs were cleaned with 70% ethanol and incubated in a poultry incubator (MultiQuip) at 37 °C, 40 - 50% humidity with constant rotation. On E4, albumen (4-5 mL) was aspirated from the acute pole of the egg using a 19G needle to generate an artificial air sac directly over the CAM, allowing its dissociation from the eggshell membrane. A circular window with a 10 mm radius was opened in the eggshell using scissors and sealed with a transparent film dressing (LivingStone) to prevent dehydration and possible infection. The eggs were returned to the incubator and incubated in a horizontal position with the window facing up and without rotation. On E8, a 5 x 5 x 2 mm diameter square gel, with or without glucose oxidase capsules, was added on top of the CAM (n = 6 per condition). PBS (50 µL) or VEGF165 (50 µL at 100 ng/mL) were added to the CAM within a 9 mm inner diameter silicone ring and served as negative and positive controls respectively. On E13, the CAM was excised and imaged under a dissecting microscope at 2.5X magnification. The number of blood vessel branches intersecting the gels were counted using ImageJ. For conditions that do not contain gels, a square with size matching the gel was drawn in the microscopic images to allow analysis consistency.

#### Tubulogenesis assays

For 2D tubulogenesis assays, Passage 5 HUVECs were used for three conditions: (1) a negative control (DMEM with 10% FBS); (2) a positive control (EGM-2 media); (3) and the capsule condition (Final normalized activity of 100 U/mL capsules). First, a thin layer of Geltrex (Thermo Fischer, A1413202) was formed on the bottom of 48 well plates (25 µL per well) on ice. Next, For the hypoxic gel conditions, 1 vol% of capsules (100 U/mL of loaded enzyme) were added to microgel suspensions which were crosslinked. These gels were then placed onto the side wall of the well for the hypoxic conditions, and the gel was slightly pushed into the uncrosslinked Geltrex. The wells were then allowed to crosslink in the incubator at 37 °C for one hour. HUVECs (20,000 cells per mL, 500µL per well) were then added to each well, and photographs were taken at 4, 8, 12, and 24 hours. The images were subsequently analyzed using the ImageJ Angiogenesis plugin.

For 3D endothelial cell tubulogenesis, HUVECS ($5 \times 10^6$ cells per mL) were seeded in microgel suspensions (medium particle, 10 wt%, 0, 0.5%, or 0.75% filler) containing 0 capsules or 2 vol% capsules with 100 U/mL of loaded glucose oxidase and 6,000 U/mL of catalase. The conditions were cultured for either 1, 5, or 7 days prior to fixation, staining with Hoechst and phalloidin, and imaging (Confocal Zeiss LSM 800, 100 μm z-stack with 2 μm slices). The images were subsequently imported into ImageJ and analyzed using the angiogenesis plugin tool. The workflow was as follows:

1. The stack had the channels split
2. A z-stack projection was made of the actin channel
3. The image was then converted to RGB
4. background was subtracted using a pixel size of 20 with a sliding palabra
5. Then contrast was enhanced with equalize histogram
6. The background was subtracted again with a ball radius 10-20 pixels
7. A gaussian blur was applied across the whole image
8. The angiogenesis tool was used on phase contrast mode.

#### Supplementary figures

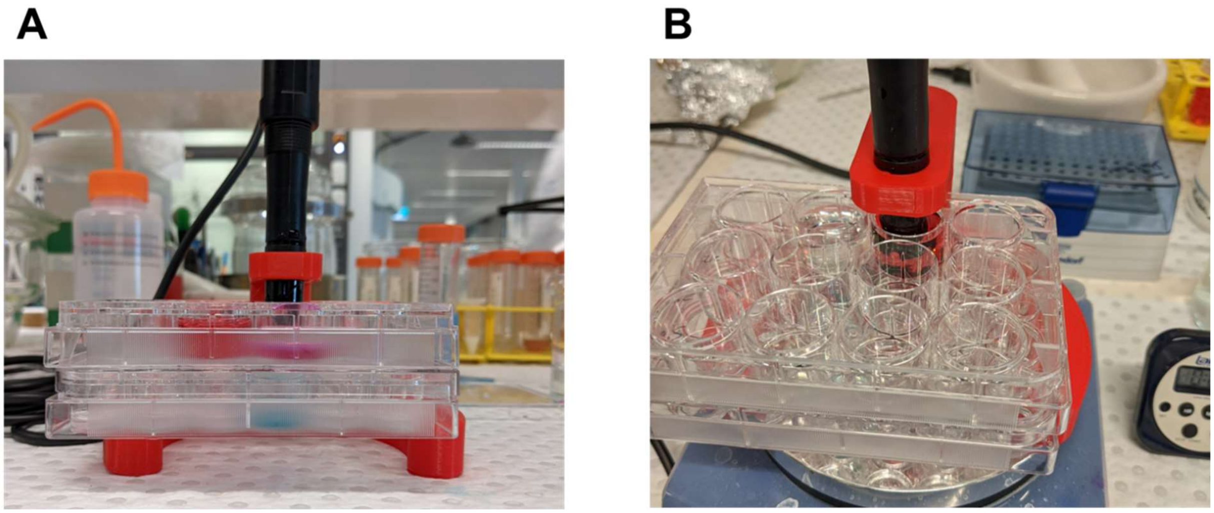

**Figure S1.** Optical images (A, B) of the Dissolve oxygen probe setup used to measure oxygen levels and enzyme kinetics.

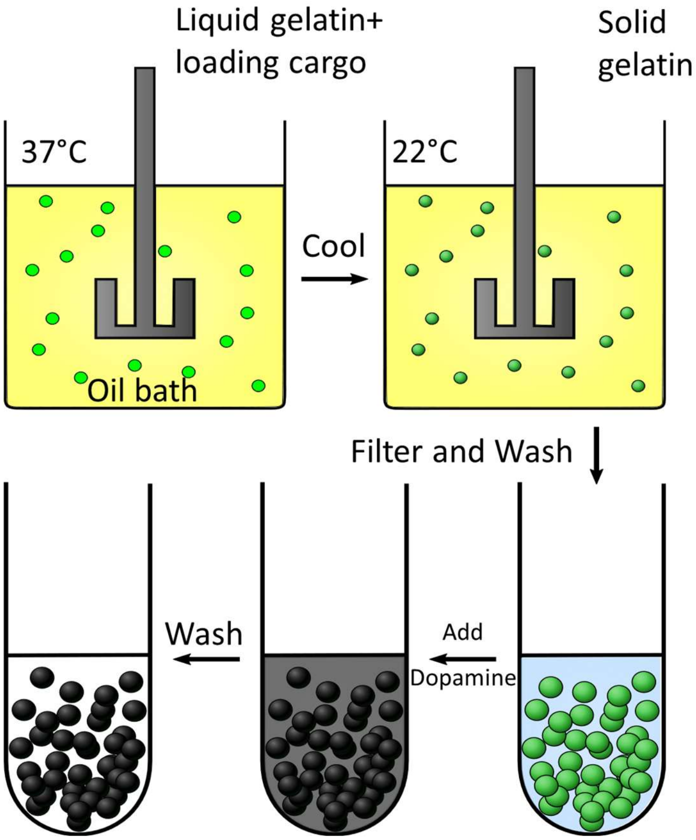


**Figure S2.** Schematic of capsule synthesis procedure. First, 37 °C gelatin is loaded into a stirred warm oil bath. After cooling and an addition of acetone, the particles are washed in hexane and air dried. Then are subsequently placed in a Tris-HCL buffer (pH 8.5 50mM) with 10 mg/mL of dopamine and shaken at 300 rpm for 24 hours before 4 PBS washes.

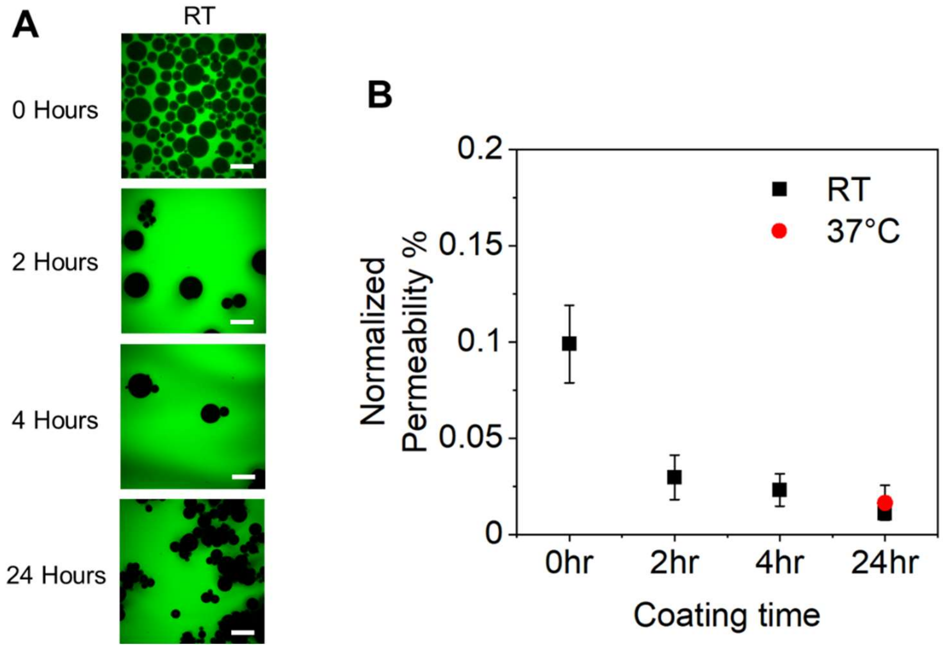


**Figure S3.** Capsule permeation characterization. A) Confocal microscopy images of capsules (coated for 0, 2, 4, or 24 hours) sitting in FITC-Dextrans for 24 hours (at room temp). B) Zoom in graph of normalized permeability of capsules based on fluorescence. Scale Bar: 100 µm.

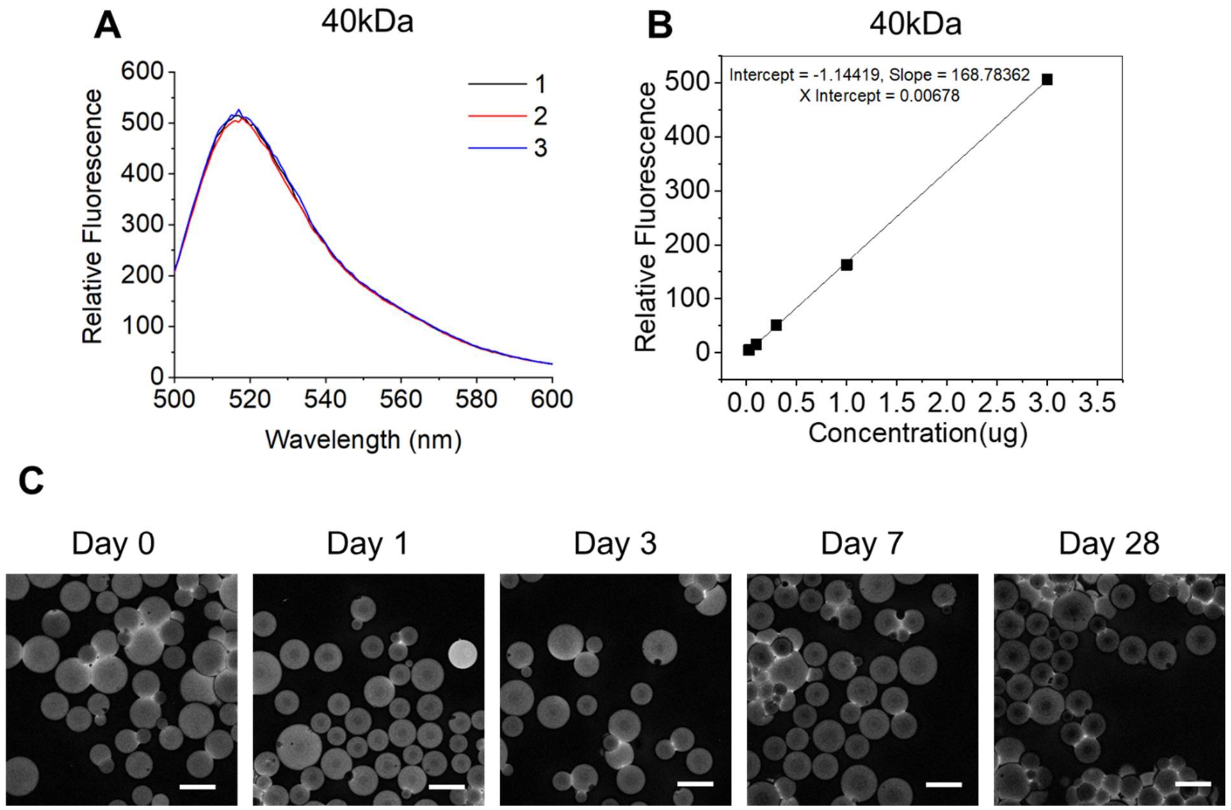


**Figure S4.** Stability of polydopamine coating. A) Spectrophotometer peaks (488 excitation) of 250kDa FITC-Dextran that has been removed from simulated-coated gelatin microgels. B) A standard curve of 250kDa FITC-Dextran used to determine concentration. C) Confocal microscopy images taken of FITC-BSA loaded microcapsules after 0, 1, 3, 7, and 28 days. Scale bars: 100 µm.

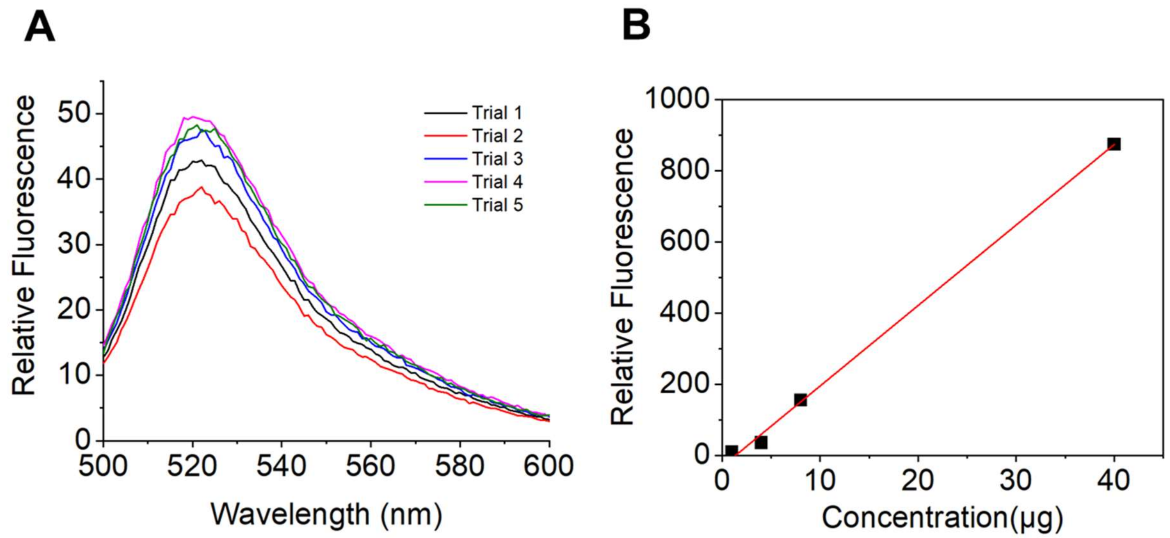


**Figure S5.** Maximal loading with the microcapsules. A) spectrophotometer curves (488 excitation) of FITC-BSA released from within "fake coated" gelatin microcapsules (100 mg/mL of BSA loaded, 1:1000 ratio BSA to FITC-BSA). B) standard curve generated for FITC-BSA on a spectrophotometer.

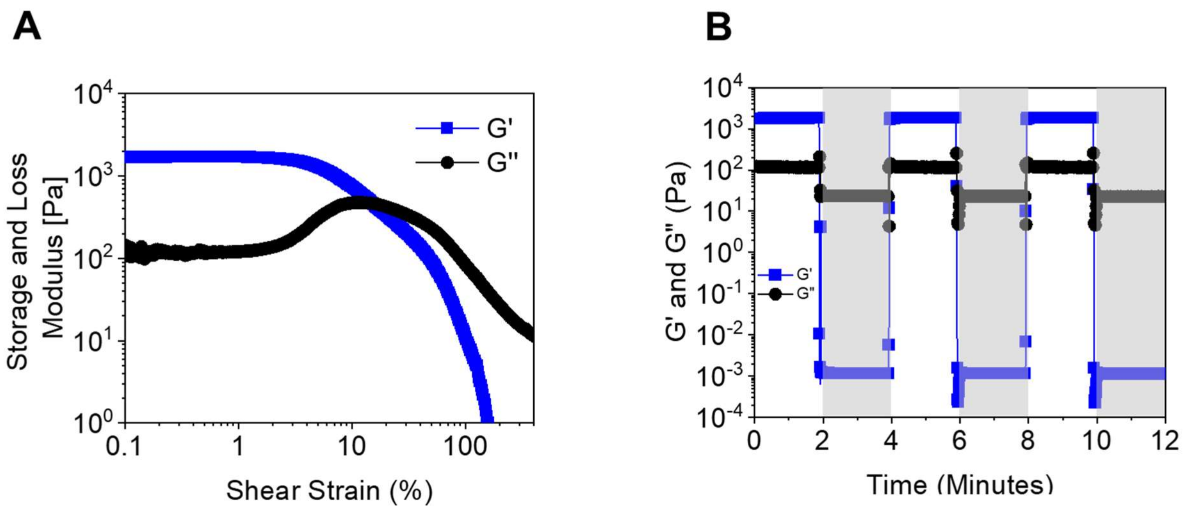


**Figure S6.** **Rheological characterizations of jammed microcapsules.** A) Shear amplitude sweeps (log ramp of 0.1-400% shear strain over 6 minutes, 1Hz). B) Thixotropic test with 3 cycles of 2 minutes of low shear strain (0.2%, 1 Hz) followed by 2 minutes at high shear strain (greyed regions) (200%, 1 Hz).

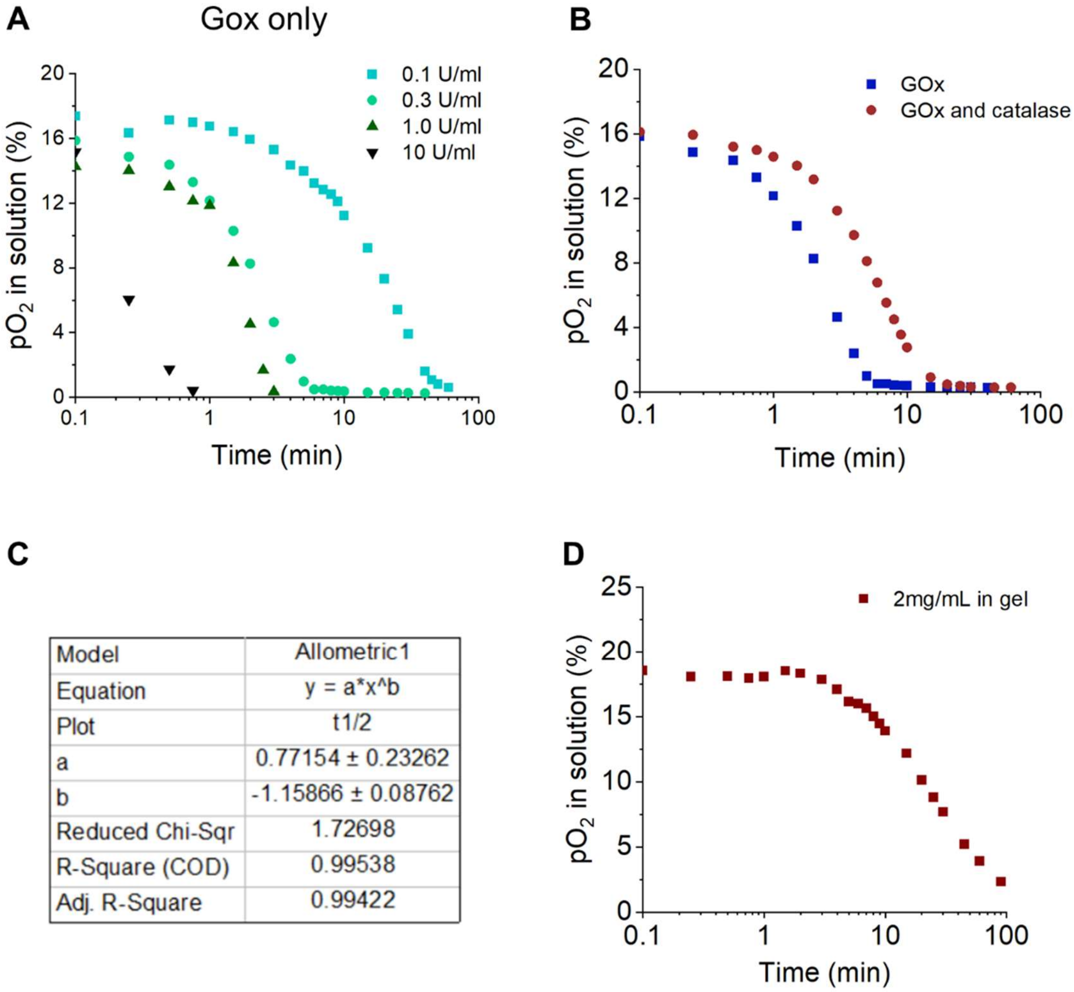


| | |
|---|---|
| Model | Allometric1 |
| Equation | y = a*x^b |
| Plot | t1/2 |
| a | 0.77154 ± 0.23262 |
| b | -1.15866 ± 0.08762 |
| Reduced Chi-Sqr | 1.72698 |
| R-Square (COD) | 0.99538 |
| Adj. R-Square | 0.99422 |

**Figure S7. Dissolved oxygen extra characterization.** A) A plot of the dissolved oxygen partial pressures for varied concentrations of glucose oxidase without catalase in cell media at room temperature. B) Partial pressures of dissolved oxygen overtime for room temperature cell media with glucose oxidase and with or without the addition of catalase. C) Data for line of best fit for saturation half time plot. D) A plot of the dissolved oxygen partial pressures for capsules loaded within a GelMa microgel suspension (20 mg/mL of glucose oxidase loaded capsules in the hydrogel, 400 µL of hydrogel in 4 mL of media (1000 U/mL, 60,000 U/mL catalase)).

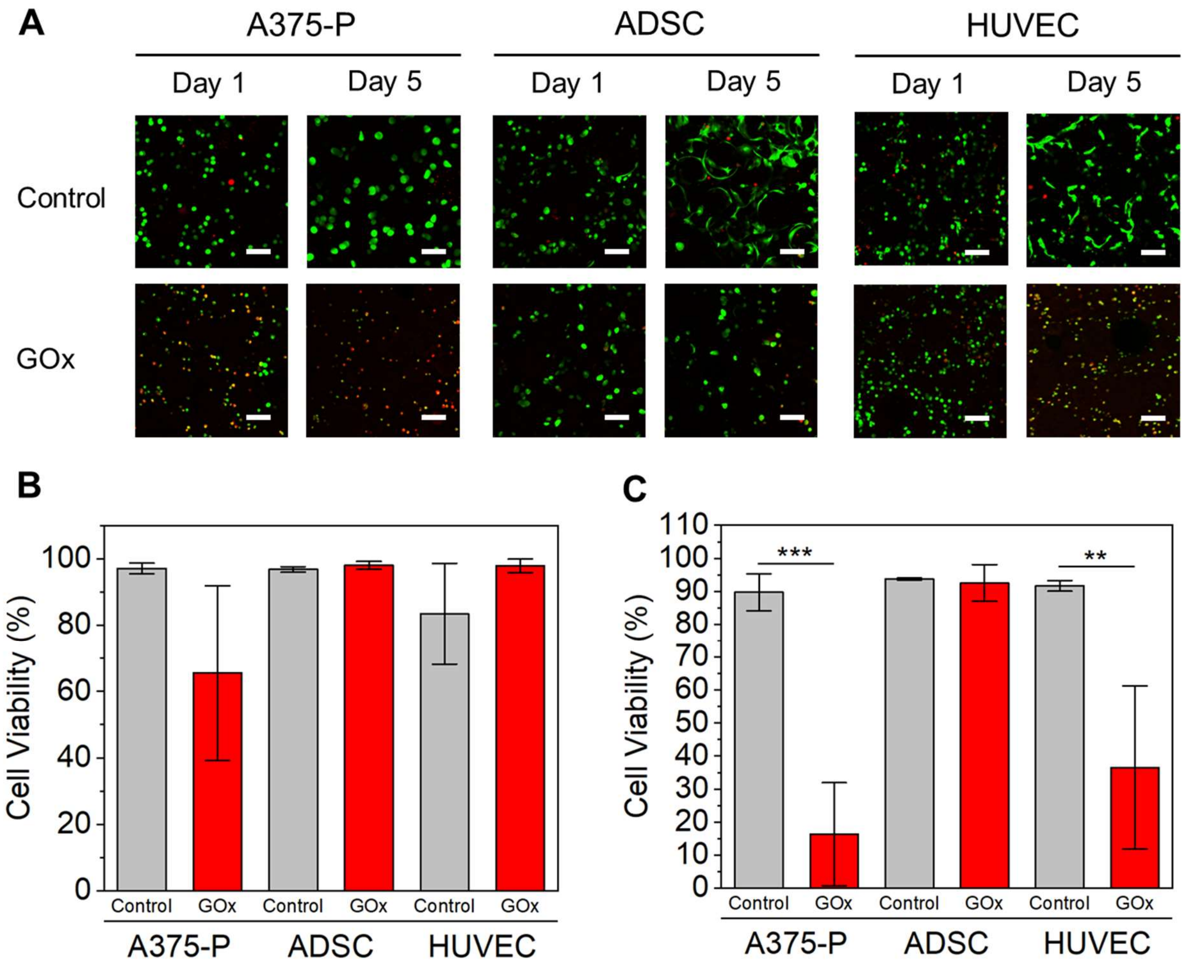


**Figure S8.** Live dead analysis for glucose oxidase loaded capsules. A) Confocal images of cells (A375-P, ADSCs, and HUVECS) stained with a live dead kit after either 1 or 5 days of culture in either control microgel suspensions (Medium, 10 wt%, 1% filler) or in suspensions with 2 vol% glucose oxidase loaded microcapsules (1000 U/mL, 60,000 U/mL catalase). Cell viability plots for all conditions at day 1 (B) and day 5 (C). Scale bars: 50 µm. ***$P < 0.001$, ANOVA. Error bars represent s.d.

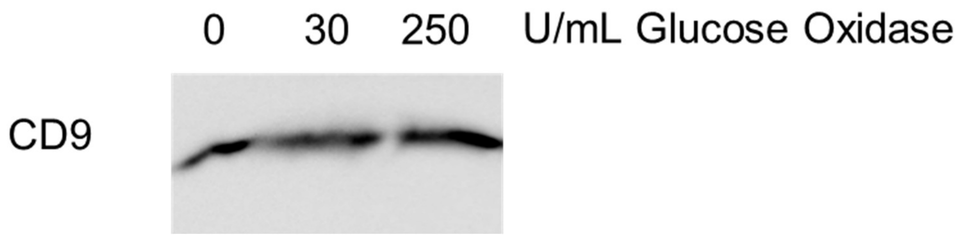


**Figure S9.** Western blot analysis showing CD9 expression in extracellular vesicles lysates (10µg) isolated from ADSCs embedded into 3D microgel suspensions with capsules containing 0, 30, or 250 U/mL of glucose oxidase.

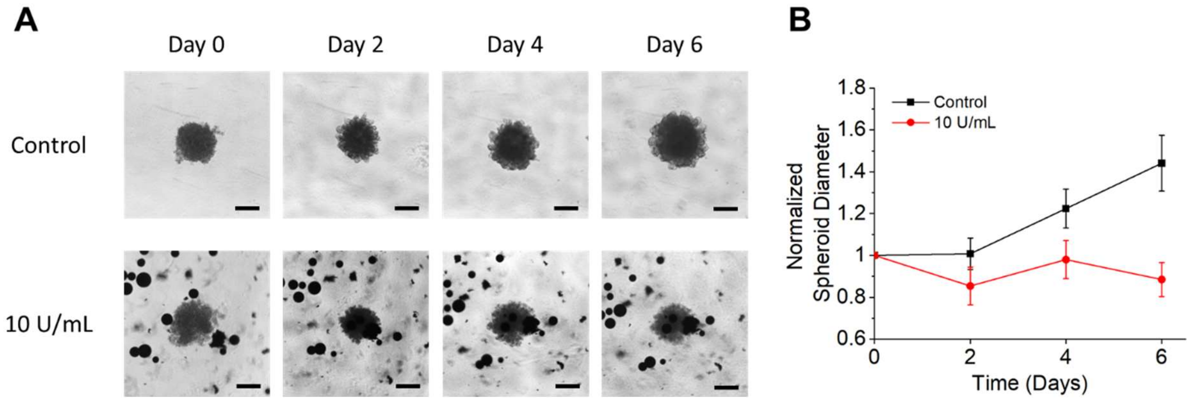


**Figure S10.** H) Optical images of MDA-MB-231 breast cancer spheroids embedded in Geltrex over 6 days with no capsules or with capsules loaded with an equivalent of 10 U/mL. I) Growth curves of breast cancer spheroids diameters. Scale bars: 200 μm.

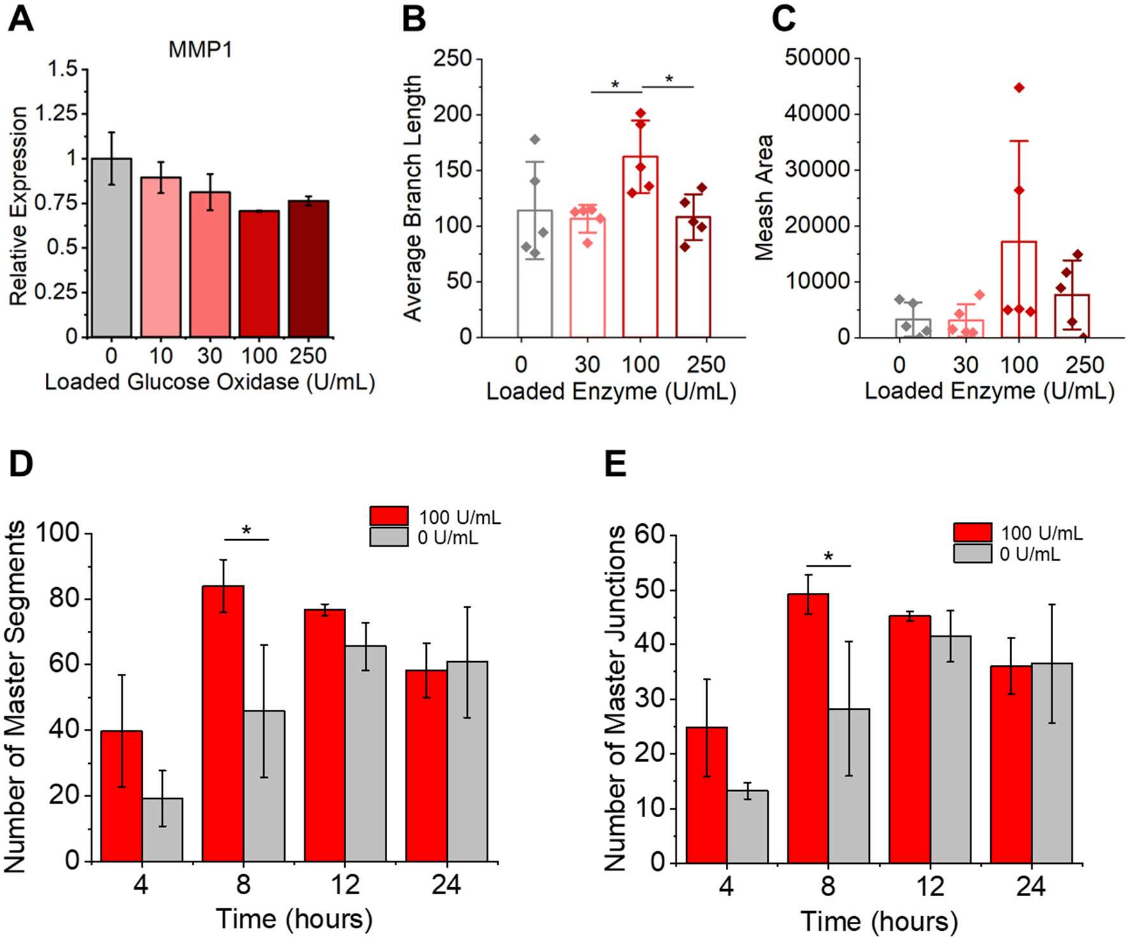


**Figure S11.** A) plots of relative expression of MMP1 for HUVECs in microgel suspensions with varied amounts of hypoxic capsules for 24 hours. Plots normalized to GAPDH. Plots of master branch length (B) and mesh area (C) and lengths for control and glucose oxidase capsule conditions calculated using the angiogenesis analyzer plugin for ImageJ. Analysis of number of master segments (D) and number of master junctions (E) from the tubulogenesis assay. *P < 0.05, ANOVA. Error bars represent s.d.

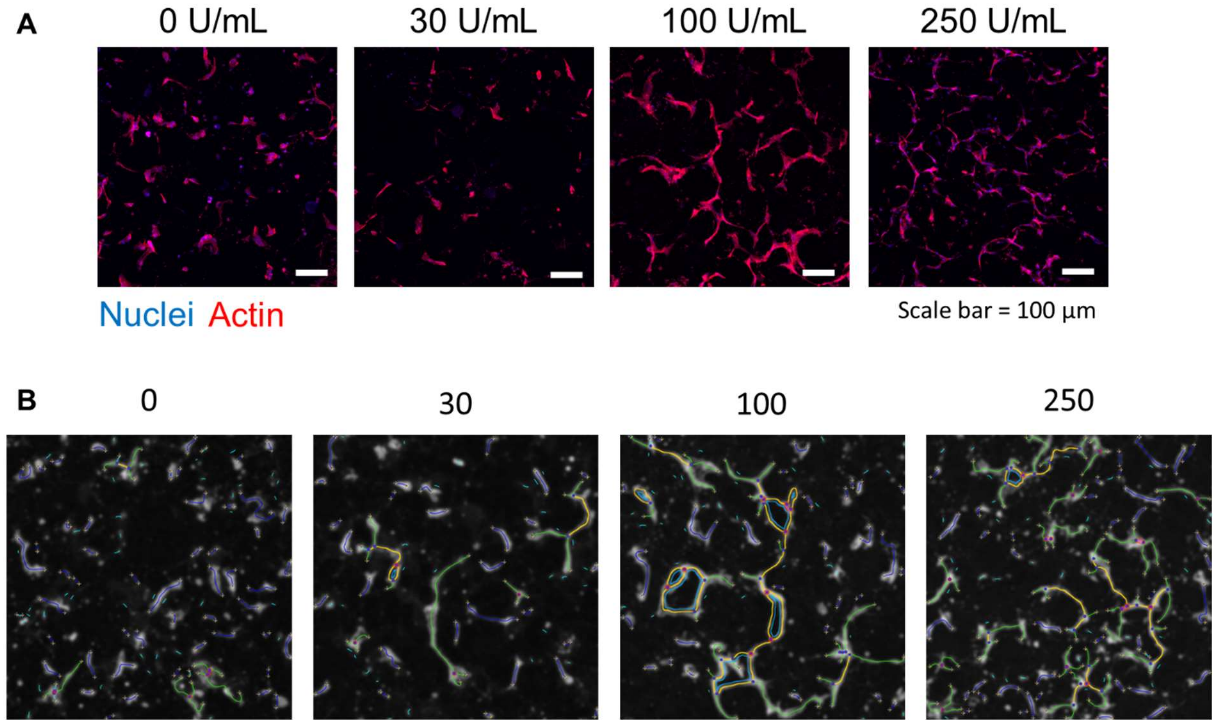


**Figure S12.** Angiogenesis in 3D microgel suspensions. A) Confocal z-stack projection images of HUVECs ($5 \times 10^6$ cells per mL, 50 slices over 100 μm z-depth) loaded in microgel suspensions with 2 vol% of capsules loaded with 0, 30, 100, or 250 U/mL of glucose oxidase enzyme. B) Outputs from the ImageJ angiogenesis analyzer tool of images of HUVECs after 5 days of culture (not the same images as from A). Scale bars: 100 μm.

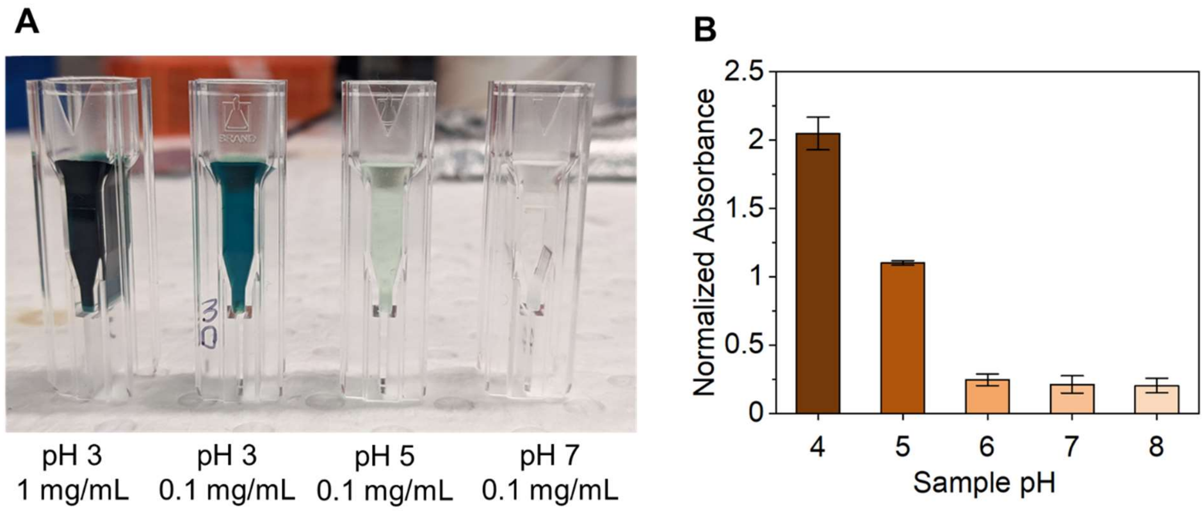


**Figure S13.** Nanoceria activity. A) Photograph of cuvettes loaded with ABTS (2mM) and nanoceria (0.1 to 1 mg/mL) in 0.1M McIlvaine buffers at various pH levels (3, 5, and 7) and left to sit overnight. B) plate reader absorbance levels of a 415 nm peak with ABTS (2mM) and nanoceria (0.5 mg/mL) in 0.1M McIlvaine buffers at various pH levels (4, 5, 6, 7, and 8) and left to sit overnight.

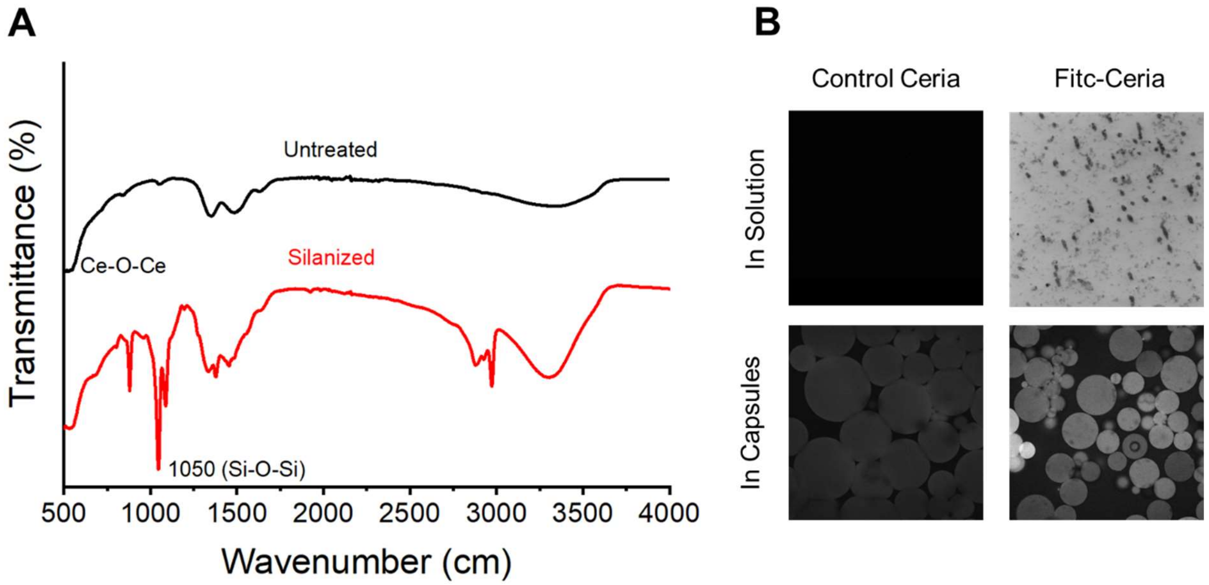


**Figure S14.** Loading fluorescent ceria in microcapsules. A) FTIR transmission plot of nanoceria with and without silane coatings. B) Confocal microscopy images of control and FITC-conjugated nanoceria particles either in solution (top) or in capsules (bottom). Scale bars: 100 µm.

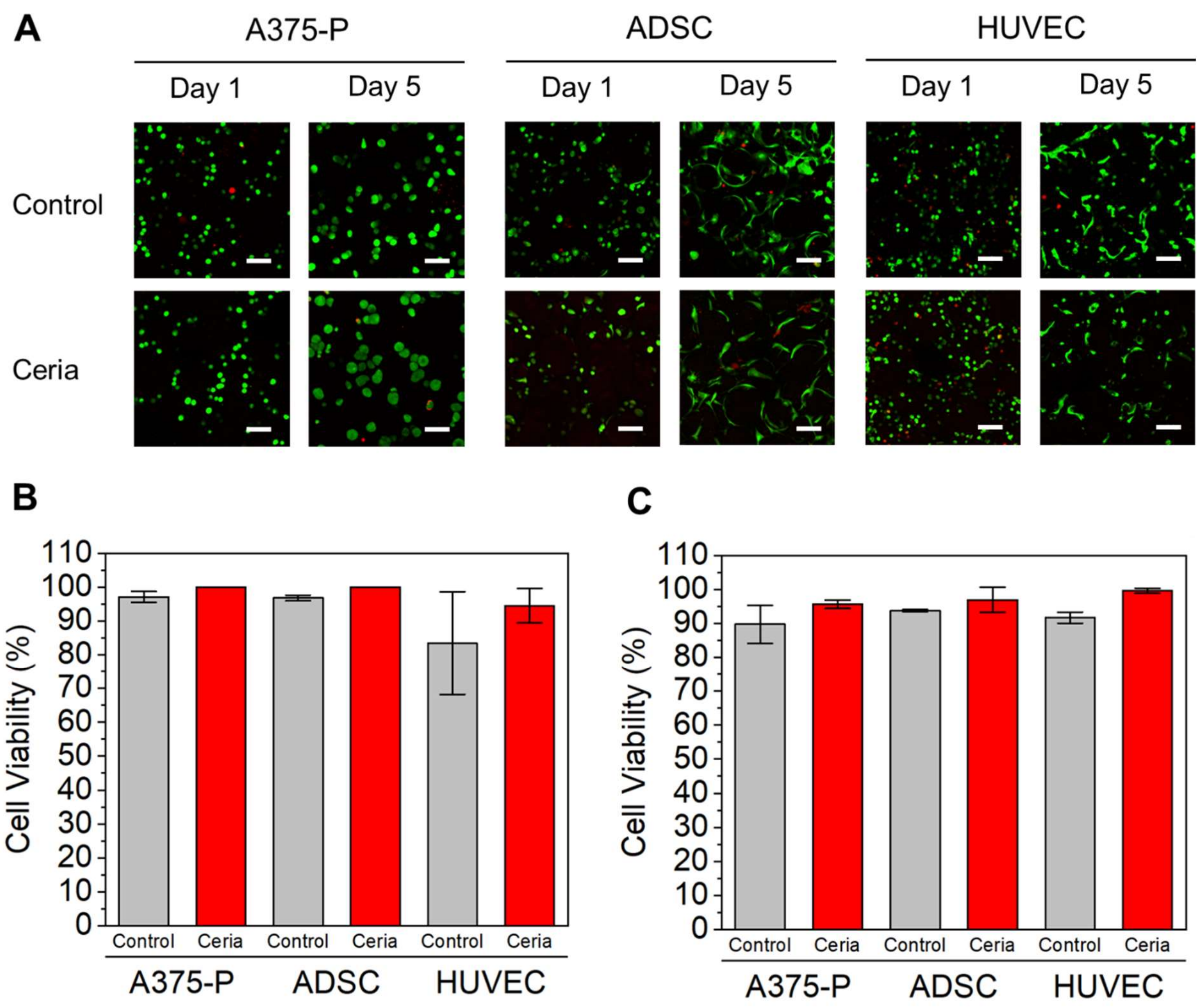


**Figure S15.** Live dead analysis for nanoceria loaded microcapsules. A) Confocal images of cells (A375-P, ADSCs, and HUVECS) stained with a live dead kit after either 1 or 5 days of culture in either control microgel suspensions (Medium, 10 wt%, 1% filler) or in suspensions with 2 vol% ceria loaded microcapsules (5 mg/mL). Cell viability plots for all conditions at day 1 (B) and day 5 (C). Scale bars: 50 μm.

**Table S1**. Media Compositions

| NAME | BASAL MEDIA | SERUM | ANTIBIOTICS/ANTIMYCOTICS | EXTRA |
| --- | --- | --- | --- | --- |
| ADSC Expansion Media | DMEM Low Glucose | 10% FBS | 1% Penicillin/Streptomycin (10,000 U/mL) | - |
| HUVEC Media | Endothelial Basal Medium | 2% FBS | Gentamicin | BulletKit: CC-3162 |
| MCF-7 MDA-MB-231 | DMEM High Glucose (4.5mg/L glucose, L-Glutamine, Sodium Pyruvate) | 10% FBS | 1% Penicillin/Streptomycin (10,000 U/mL) | |
| A375-P | DMEM High Glucose | 10% FBS | 1% Penicillin/Streptomycin (10,000 U/mL) | 1% Glutamax |

**Table S2**. Immunofluorescent stains

| Stain | Company | Catalog # | Dilution |
|---|---|---|---|
| Hoechst 33342 | Life Technologies Australia Pty ltd | 687117 | 1:200 |
| Phalloidin-Atto 488 | Sigma-Aldrich | 49409 | 1:100 |
| Phalloidin-Atto 647 | Sigma-Aldrich | 65906 | 1:250 |
| Anti-HIF-1 alpha antibody [EP1215Y | Abcam | ab51608 | 1:250 |
| Goat Anti-Rabbit IGG 488 | Life Technologies Australia Pty ltd | 35553 | 1:250 |
| CD9 (D8O1A) Rabbit | Cell Signaling Technology | 13174S | 1:1000 |
| Anti-rabbit IgG, HRP-linked Antibody | Cell Signaling Technology | 7074P2 | 1:5000 |
| Calcein AM | Life Technologies Australia Pty ltd | L3224 | 2 µM |
| Ethidium Homodimer-1 | Life Technologies Australia Pty ltd | L3224 | 4 µM |
